# Proteostasis Remodeling Across Development Defines Fetal, Neonatal, and Adult Hematopoietic Stem Cell States

**DOI:** 10.64898/2025.12.12.694052

**Authors:** Helena Yu, Yoon Joon Kim, Katelyn Chen, Andrea Z. Liu, Mary Jean Sunshine, Robert A.J. Signer

## Abstract

Hematopoietic stem cells (HSCs) must preserve protein homeostasis (proteostasis) despite dramatic changes in proliferative and biosynthetic demands during development, yet how proteostasis is regulated across these transitions is poorly understood. Here, we show that fetal and neonatal HSCs operate through distinct, stage-specific proteostasis programs that differ fundamentally from those in adulthood. Using quantitative in vivo assays spanning embryonic through adult stages, we uncover an unanticipated decoupling between protein synthesis and protein quality control during development, revealing that fetal and neonatal HSCs employ specialized mechanisms to safeguard proteome integrity under developmental stress. Developing HSCs experience a distinctive proteostasis landscape characterized by elevated protein synthesis, increased unfolded protein burden, and selective engagement of stress-buffering and protein degradation pathways that are largely dispensable in young adult HSCs. Disruption of these pathways compromises early life HSC function and long-term fitness, establishing proteostasis control as a key regulator of stem cell maturation. These findings define previously unrecognized mechanisms by which HSCs manage the proteome during early life and reveal fundamental principles governing stem cell proteostasis across ontogeny.

## INTRODUCTION

Hematopoietic stem cells (HSCs) must maintain an exceptionally pristine proteome to sustain lifelong regenerative capacity. Protein homeostasis (proteostasis) - the coordinated regulation of protein synthesis, folding, trafficking, and degradation - is therefore a fundamental determinant of HSC integrity^1–16^. In young adult HSCs, low protein synthesis^2^ and stringent quality control pathways preserve proteome integrity^3,4^, and even subtle disruption of these processes impairs HSC self-renewal, fitness, and long-term function^2–4^. Despite this central importance, how the proteostasis network is organized in HSCs during early life remains largely unexplored.

Ontogeny introduces profound shifts in HSC physiology, including changes in cell cycle behavior^17–22^, metabolic wiring^15,23–27^, lineage output^28–37^, and stress tolerance^5,16,38–43^. Prior work indicates that fetal HSCs sustain substantially higher protein biosynthetic loads and operate in distinct metabolic states and environments compared with their adult counterparts^25,38,44,45^.

These developmental transitions likely reshape the demands placed on proteostasis machinery, but the architecture of proteostasis in developing HSCs has never been systematically defined. In particular, how protein synthesis rates, misfolded protein burden, and engagement of degradation pathways change across fetal, neonatal, and young adult HSCs remains unknown.

This gap is notable because developmental stage is increasingly recognized as a major modifier of HSC vulnerability to dysfunction and disease^44,46–56^. Pediatric blood disorders ranging from congenital bone marrow failure syndromes to childhood leukemias often arise in a developmental context in which proteostasis regulation and stress may be distinct from adults^44,55^. Without a foundational map of how proteostasis pathways operate during early life in physiologic development, it remains difficult to determine whether disease-associated mutations intersect uniquely with fetal and neonatal proteostasis demands, or whether developmental windows expose HSCs to vulnerabilities that are not evident in adulthood.

Here, we address this unmet need by defining proteostasis control across key stages of HSC ontogeny. We mapped developmental changes in protein synthesis, unfolded and misfolded protein burden, and activity of major protein degradation pathways, and determined how perturbations to fetal proteostasis pathways impact HSC function and long-term hematopoietic fitness. These analyses uncovered unanticipated principles of how developing HSCs maintain proteome integrity and reveal developmental windows in which proteostasis becomes uniquely strained. This framework establishes a foundation for understanding how early-life perturbations in proteostasis may influence pediatric hematologic disease.

## RESULTS

### Protein synthesis rates progressively decline in HSCs after birth

Low protein synthesis is a defining feature of adult HSCs and is essential for maintaining proteostasis and long-term self-renewal^2^. We previously showed that fetal HSCs synthesize protein at far higher rates than adult HSCs^44^, but whether the transition from high to low protein synthesis occurs gradually or abruptly remained unclear. To address this, we performed time-mated analysis and quantified protein synthesis at 6 developmental stages in vivo (E16.5, E18.5, P0, P7, P14, 8-weeks-old, Fig. 1A). Pregnant dams, neonates, and young adult mice were injected with O-propargyl-puromycin (OP-Puro)^2,57,58^, and incorporation was measured 1h later by flow cytometry in CD150^+^CD48^-^Lineage^-^Sca1^+^ckit^+^ (CD150^+^CD48^-^LSK) HSCs^59^.

**Figure 1.**
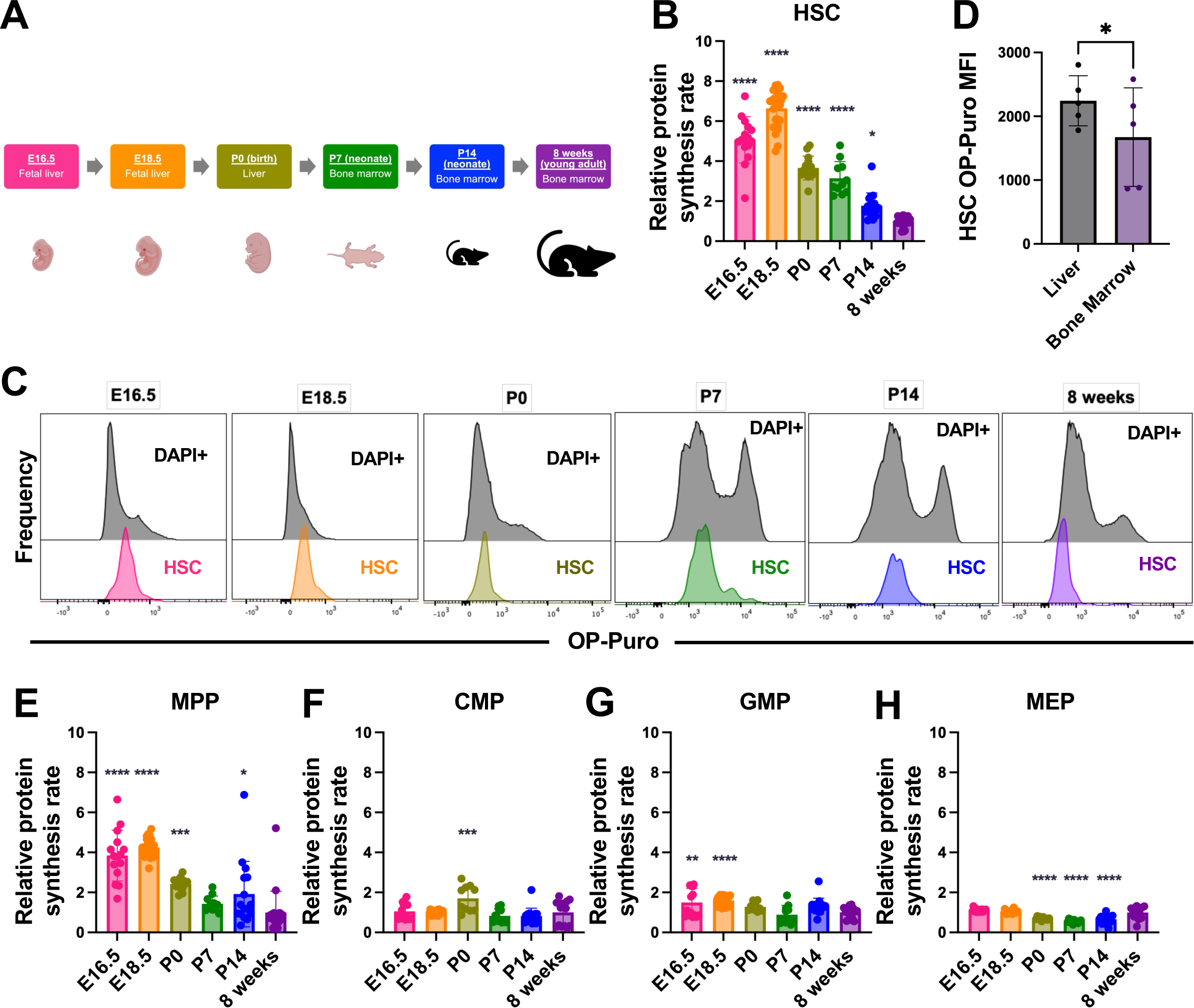
Protein synthesis rates progressively decline in HSCs after birth (A) Schematic showing ages of mice analyzed and the tissue source of hematopoietic stem and progenitor cells. (B) Relative protein synthesis rates in HSCs at E16.5, E18.5, P0, P7, and P14 as measured by OP-Puro mean fluorescence intensity (MFI) normalized to unfractionated DAPI^+^ cells, compared to normalized OP-Puro MFI of HSCs from 8-week-old mice (n = 11-20 mice/age in 2-4 experiments/age). (C) Representative histograms of OP-Puro of DAPI^+^ and HSC populations at E16.5, E18.5, P0, P7, P14, and 8 weeks. (D) OP-Puro MFI in P0 liver and bone marrow HSCs (n = 5 mice). (E-H) Relative protein synthesis rates in MPPs (E), CMPs (F), GMPs (G), and MEPs (G) at E16.5, E18.5, P0, P7, and P14 as measured by OP-Puro MFI normalized to unfractionated DAPI^+^ cells, compared to normalized OP-Puro MFI from 8-week-old mice (n = 10-20 mice/age in 2-4 experiments/age). Data represent mean ± standard deviation (B, E-H). Statistical significance was assessed using a one-way ANOVA followed by Dunnett’s multiple comparisons test relative to the 8-week-old timepoint. *P<0.05; **P<0.01; ***P<0.001; ****P<0.0001.

OP-Puro incorporation was highest during fetal development and peaked at E18.5, reaching 6.6-fold greater levels than in young adult (8-week-old) HSCs when normalized to unfractionated cells (Fig. 1B; P<0.0001). Fetal HSCs exhibited protein synthesis rates among the highest of all cells in the fetal liver (Fig. 1C). After birth, HSC protein synthesis declined steadily yet remained significantly elevated relative to adult HSCs at every postnatal stage examined (Fig. 1B). P0 and P7 HSCs showed 3.7 and 3.1-fold higher protein synthesis than adults (P<0.0001), respectively, whereas P14 HSCs maintained 1.8-fold higher rates (Fig. 1B; P<0.05). At these postnatal timepoints, HSCs exhibited intermediate protein synthesis compared with unfractionated cells, whereas adult HSCs had amongst the lowest protein synthesis rates (Fig. 1C). OP-Puro incorporation in HSCs displayed largely unimodal distributions across all stages, with only minor broadening at P7 (Fig. 1C). These data indicate that developmental shifts in protein synthesis occur uniformly across the HSC compartment rather than through emergence of asynchronous or transient HSC subsets.

Because the hematopoietic niche transitions from fetal liver to bone marrow^60–64^, we tested whether this shift contributes to reduced protein synthesis. Paired analysis of P0 liver and bone marrow revealed a modest but significant reduction in OP-Puro incorporation in bone marrow HSCs (Fig. 1D). Nevertheless, P7 and P14 bone marrow HSCs all maintained significantly higher protein synthesis than adult HSCs (Fig. 1B), suggesting that the bone marrow environment may contribute partially but cannot fully account for the postnatal decline. These findings imply that niche signals act in concert with intrinsic developmental programs to progressively constrain protein synthesis.

Differences in protein synthesis also cannot be explained solely by variation in cell cycle activity. Prior work has shown that forcing adult HSCs into rapid self-renewing divisions in vivo increases protein synthesis by only ∼2-fold^2^, which is far below the elevation in protein synthesis observed in fetal HSCs (Fig. 1B).

To determine whether this developmental pattern is specific to HSCs or a broader feature of hematopoiesis, we quantified OP-Puro incorporation across CD150^-^CD48^-^LSK multipotent progenitors^65^ (MPPs) and several restricted myeloid progenitor populations^66^ (CMPs, GMPs and MEPs). Adult MPPs exhibited protein synthesis rates comparable to adult HSCs, and their developmental trajectory of translational activity was similar but less pronounced. MPP protein synthesis peaked during fetal life at 4.3-fold above adult levels and then declined more rapidly than HSCs after birth, with P7 MPPs exhibiting protein synthesis rates that were only modestly elevated and not significantly different from adults (Fig. 1E).

Restricted progenitors showed minimal dynamic changes in protein synthesis during development. CMPs exhibited largely constant protein synthesis across ontogeny, with only a modest increase at P0 (Fig. 1F). GMPs displayed 1.6-fold elevation in protein synthesis during fetal development, with all postnatal stages resembling adults (Fig. 1G). MEPs showed reduced protein synthesis at P0, P7, and P14 relative to adults (Fig. 1H). Collectively, the magnitude of developmental changes in protein synthesis within restricted progenitors was far smaller than in HSCs. Thus, the dramatic modulation of protein synthesis is uniquely concentrated within the stem cell compartment.

Together, these data demonstrate that protein synthesis is tightly and dynamically regulated in HSCs across development: extremely high during fetal life, progressively declining after birth, and reaching exceptionally low levels in young adults.

### Developing HSCs accumulate unfolded and misfolded protein

Adult HSCs maintain proteostasis in part by restricting global protein synthesis^2^, thereby limiting the biogenesis and burden of unfolded and misfolded proteins^3^. This raised the question of whether developing HSCs, which synthesize protein at substantially higher rates (Fig. 1B), accumulate greater levels of aberrant proteins. To test this, we quantified unfolded and misfolded proteins in HSCs using tetraphenylethylene maleimide (TMI) fluorescence and ubiquitylated protein abundance, respectively. TMI is a cell-permeable dye that fluoresces upon binding to free cysteine thiols, which are predominantly exposed in unfolded proteins^67^.

Polyubiquitylated proteins serve as a well-established surrogate for misfolded protein load^68^. We previously validated both assays in HSCs^3^, and they have since been widely adopted by the field^4,6,55,69–72^.

Fetal HSCs at E16.5 and E18.5 exhibited ∼2-fold higher levels of unfolded and misfolded protein than adult HSCs (Fig. 2A-D). Neonatal HSCs also exhibited significantly increased ubiquitylated protein compared with adults, although TMI fluorescence was not significantly elevated at these stages (Fig. 2A,C). Despite these increases, the magnitude of aberrant protein accumulation was far more modest than expected given the large developmental differences in protein synthesis rates. Prior work suggested a more linear relationship between protein synthesis rates and unfolded protein burden in HSCs^3^, yet misfolded protein abundance varied only modestly across ontogeny. Thus, although fetal and neonatal HSCs synthesize protein at markedly higher rates, the accompanying increase in misfolded protein is proportionally modest, suggesting the presence of compensatory proteostasis mechanisms that buffer early developmental protein biosynthetic load. Similar trends were observed across progenitor populations. Developing progenitors contained higher unfolded and/or misfolded proteins (Fig. 2E-L) despite relatively little variation in their protein synthesis rates throughout development (Fig. 1F-H).

**Figure 2.**
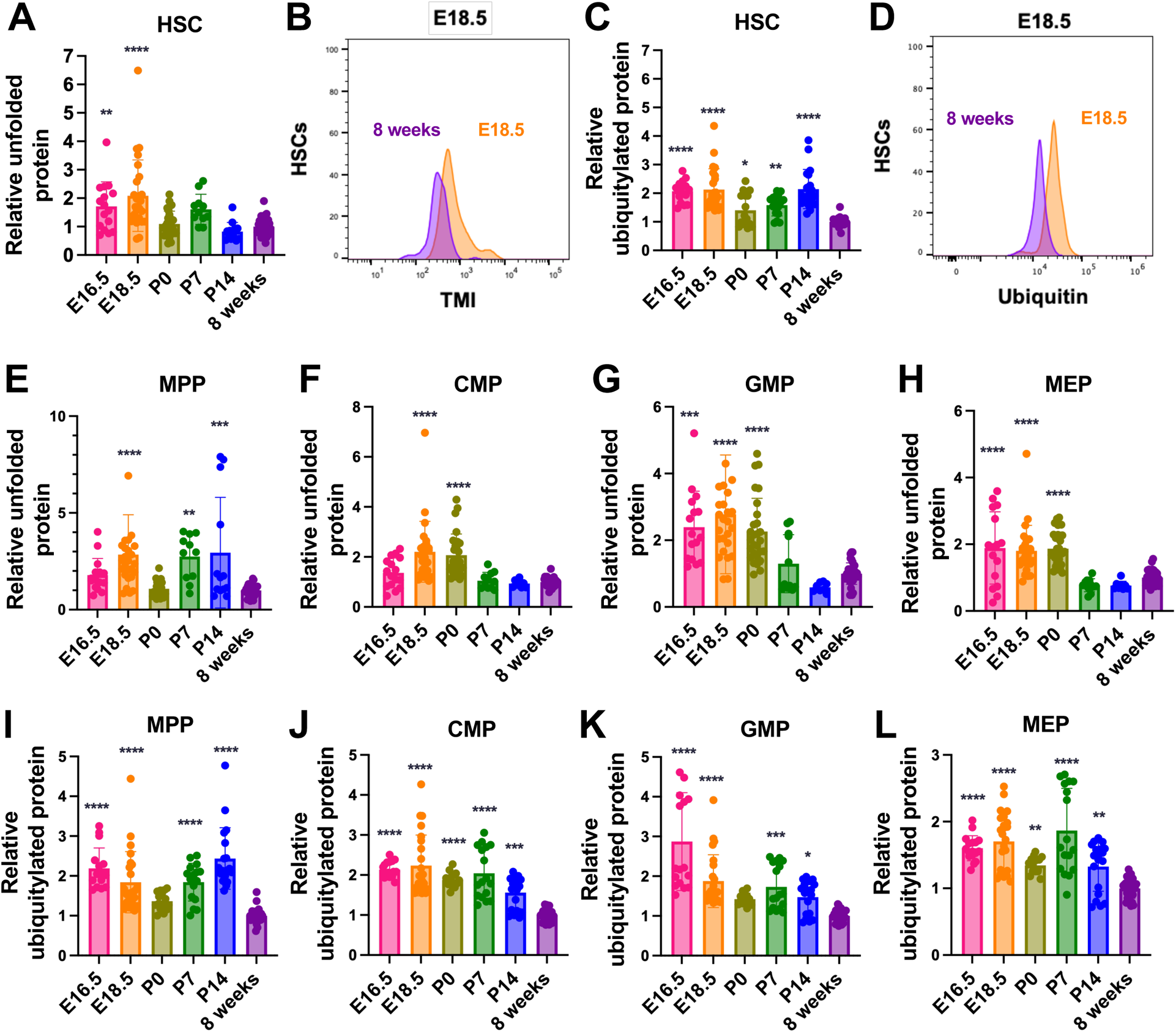
**Developing HSCs accumulate unfolded and misfolded protein.** (A) Relative quantity of unfolded protein in HSCs at E16.5, E18.5, P0, P7, and P14 as measured by tetraphenylethene maleimide (TMI) MFI normalized to unfractionated cells, compared to normalized TMI MFI of HSCs from 8-week-old mice (n = 11-37 mice/age in 2-4 experiments/age). (B) Representative histogram of TMI in HSCs at E18.5 and 8 weeks. (C) Relative quantity of ubiquitylated protein in HSCs normalized to unfractionated DAPI^+^ cells at E16.5, E18.5, P0, P7, and P14 compared to normalized values for HSCs from 8-week-old mice (n = 16-34 mice/age in 2-3 experiments/age). (D) Representative histogram of ubiquitylated protein in HSCs at E18.5 and 8 weeks. (E-H) Relative quantity of unfolded protein in MPPs (E), CMPs (F), GMPs (G), and MEPs (H) at E16.5, E18.5, P0, P7, and P14 as measured by TMI MFI normalized to unfractionated cells, compared to normalized TMI MFI of 8-week-old mice (n = 11-37 mice/age in 2-4 experiments/age). (I-L) Relative quantity of ubiquitylated protein in MPPs (I), CMPs (J), GMPs (K), and MEPs (L) normalized to unfractionated DAPI^+^ cells at E16.5, E18.5, P0, P7, and P14 compared to normalized values for 8-week-old mice (n = 16-34 mice/age in 2-3 experiments/age). Data represent mean ± standard deviation (A, C, E-L). Statistical significance was assessed using a one-way ANOVA followed by Dunnett’s multiple comparisons test relative to the 8-week-old timepoint. *P<0.05; **P<0.01; ***P<0.001; ****P<0.0001.

These findings indicate that developing HSCs carry a higher burden of unfolded and/or misfolded protein than adult HSCs, revealing a potential state of proteostasis stress.

Importantly, the modest magnitude of this increase relative to their markedly elevated protein synthesis suggests a partial uncoupling between protein production and protein quality control during development, indicating that additional proteostasis pathways must be engaged to at least partly buffer the high protein biosynthetic load.

### Neonatal HSCs have increased dependence on the proteasome to maintain proteostasis

To identify pathways that help developing HSCs limit the accumulation of unfolded and misfolded protein, we assessed protein degradation mechanisms across ontogeny. We first analyzed published RNA-sequencing datasets (GSE128759)^73^ to examine developmental changes in gene sets related to protein degradation. We recently reported that, unlike most cells that rely primarily on the ubiquitin proteasome system^74^, adult HSCs rely heavily on autophagy to clear aberrant proteins^4^. Consistent with this, expression of the “Positive Regulation of Macroautophagy” gene set remained relatively constant across all HSC developmental stages relative to adults (Fig. 3A-E), and autophagic activity was high in all HSC developmental stages (Supplementary Fig. S1A-G).

**Figure 3.**
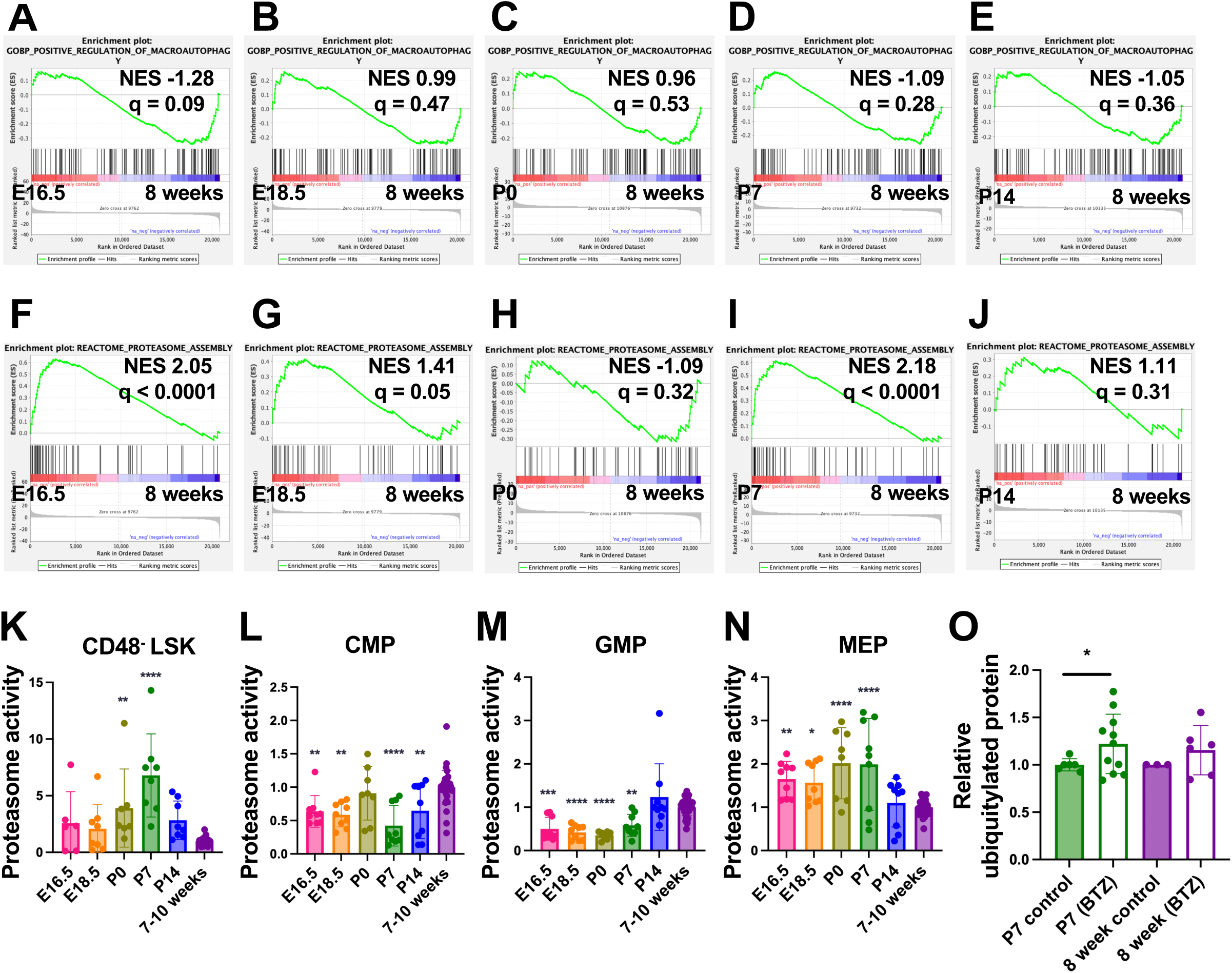
**Neonatal HSCs have increased dependence on the proteasome to maintain proteostasis** (A-E) Enrichment plots for the “Positive Regulation of Macroautophagy” gene set based on RNA-sequencing data (GSE128759) in HSCs^73^ from E16.5 (A), E18.5 (B), P0 (C), P7 (D), and P14 (E) mice compared to 8-week-old mice. (F-J) Enrichment plots for the “Proteasome Assembly” gene set based on RNA-sequencing data in HSCs from E16.5 (F), E18.5 (G), P0 (H), P7 (I), and P14 (J) mice compared to 8-week-old mice. (K-N) Proteasome activity as measured by Proteasome-Glo assay in CD48^-^LSK HSCs/MPPs (K), CMPs (L), GMPs (M), and MEPs (N) at E16.5, E18.5, P0, P7, and P14, relative to young adult (7–10-week-old) mice (n = 6-32 sets of 5000 cells from 12-27 mice/age in 3 experiments/age). (O) Relative ubiquitylated protein in HSCs from P7 and 8-week-old mice after injection with 10 mg/kg bortezomib (BTZ) as compared to non-treated age-matched controls (n = 6-11 mice/treatment group in 3-4 experiments/age). Data represent mean ± standard deviation (K-O). Normalized enrichment scores (NES) and false discovery rates (q) are shown in A-J. Statistical significance was assessed using a one-way ANOVA followed by Dunnett’s multiple comparisons test relative to the 7-10-week-old timepoint (K-N) or Welch’s t-tests (O). *P<0.05; **P<0.01; ***P<0.001; ****P<0.0001.

In contrast, several developing HSC populations, including E16.5 and P7, showed significant upregulation of the “Proteasome Assembly” gene set compared with adult HSCs (Fig. 3F-J), suggesting that the proteasome may be more active and more essential during development. To test this possibility, we quantified proteasome activity in HSCs and progenitors. Because fetal HSCs cannot be reliably assayed using this approach due to limiting cell numbers, we analyzed CD48^-^LSK HSCs/MPPs. Most developing HSCs/MPPs exhibited higher proteasome activity than adults, with P7 HSCs displaying a particularly striking 6.8-fold increase (Fig. 3K; P<0.0001). As with developmental changes in protein synthesis, this pattern was largely confined to HSCs and not observed in restricted progenitors (Fig. 3L-N). These data, together with the relatively restrained accumulation of aberrant proteins in developing HSCs, suggest that enhanced proteasome function may serve as one component of a broader developmental buffering system that maintains proteostasis despite elevated biosynthetic load.

To determine whether neonatal HSCs functionally depend on their elevated proteasome activity to maintain proteostasis, we treated P7 and young adult mice with the proteasome inhibitor bortezomib and assessed ubiquitylated protein accumulation. In agreement with prior findings showing limited proteasome contribution to HSC proteostasis in adults^4^, bortezomib treatment did not significantly increase ubiquitylated protein in young adult HSCs (Fig. 3O). In sharp contrast, P7 HSCs accumulated significant ubiquitylated protein following proteasome inhibition (Fig. 3O). Together, these data show that neonatal HSCs have elevated proteasome activity and rely more heavily on the proteasome to maintain proteostasis, reinforcing the idea that distinct proteostasis architectures operate at different developmental stages.

### Fetal HSCs activate Hsf1

To further define how developing HSCs maintain proteostasis and respond to elevated baseline levels of unfolded and misfolded protein, we assessed activation of proteostasis stress response pathways. Gene signatures related to the unfolded protein response (UPR), including “Regulation of ER UPR” and “PERK mediated UPR” showed no significant enrichment in developing HSCs (Supplementary Fig. S2). In contrast, gene set enrichment analysis revealed significantly elevated expression of the “Regulation of HSF1 Mediated Heat Shock Response” signature in E16.5 HSCs (Fig. 4A). This pathway is the principal cytosolic proteostasis stress response and is controlled by the transcription factor Heat shock factor 1 (Hsf1)^75,76^. We previously showed that Hsf1 is largely inactive in young adult HSCs, becoming activated under proteostasis stress in vitro or in response to stress associated with physiological aging^77^.

**Figure 4.**
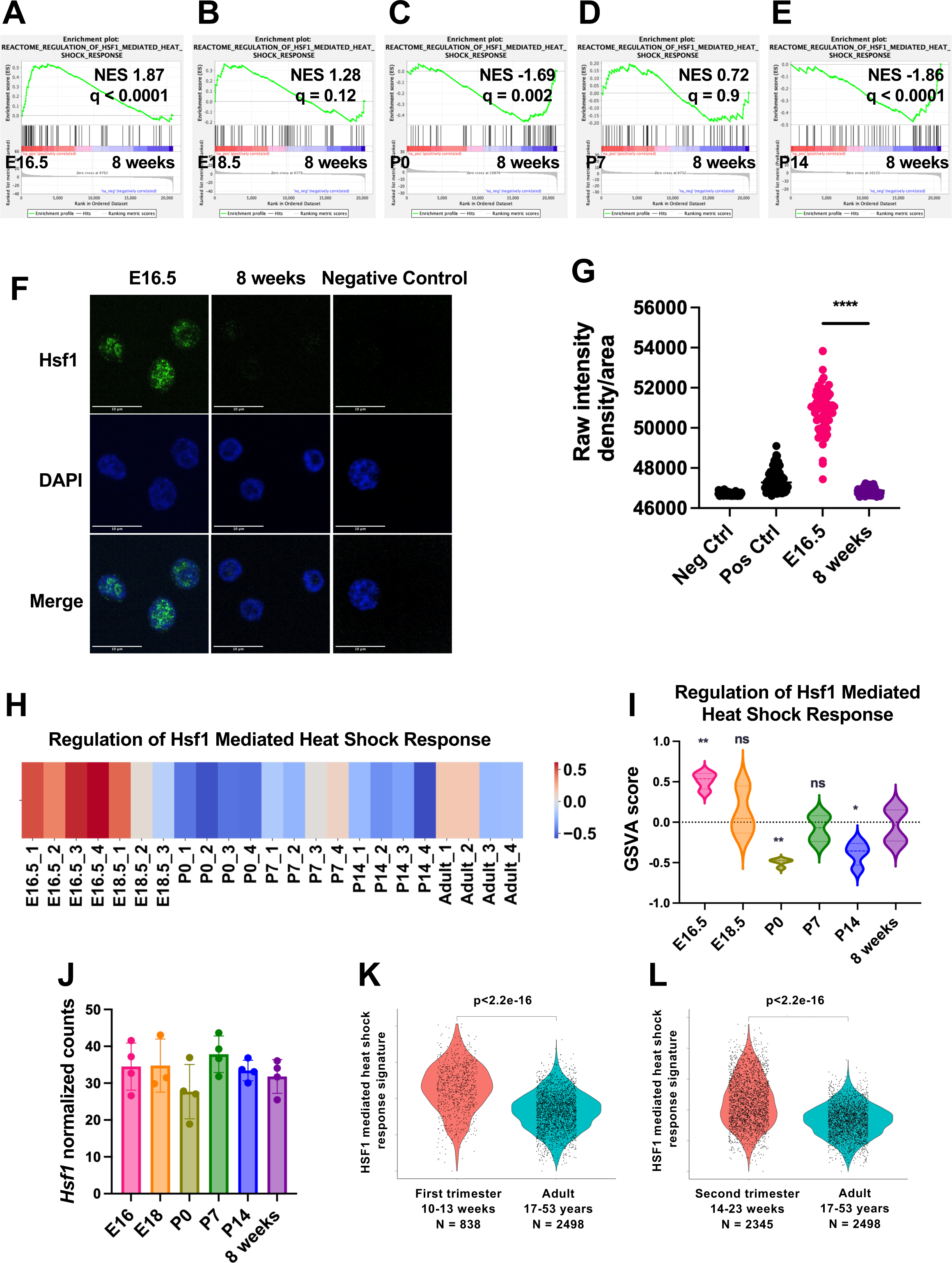
**Fetal HSCs activate Hsf1** (A-E) Enrichment plots for the “Regulation of Hsf1 mediated heat shock response” gene set based on RNA-sequencing data (GSE128759) in HSCs from E16.5 (A), E18.5 (B), P0 (C), P7 (D), and P14 (E) mice compared to 8-week-old mice. (F) Representative images of Hsf1 immunofluorescence staining in E16.5 and 8-week-old HSCs. (G) Quantification of nuclear Hsf1 immunofluorescence in E16.5 and 8-week-old HSCs (n = 72-75 HSCs). (H-I) Heat map (H) and violin plot (I) showing gene set variation analysis scores for expression of the “Regulation of Hsf1 mediated heat shock response” gene set in HSCs at E16.5, E18.5, P0, P7, P14, and 8 weeks (n = 3-4 per age). (J) Normalized counts for *Hsf1* expression in HSCs based on RNA-sequencing at E16.5, E18.5, P0, P7, P14, and 8 weeks (n = 3-4 per age). (K-L) Violin plots comparing expression of the “HSF1 mediated heat shock response” gene set in human (K) first trimester and (L) second trimester HSPCs compared to adult HSPCs. Data represent mean ± standard deviation (I). Normalized enrichment scores (NES) and false discovery rates (q) are shown in A-E. Statistical significance was assessed using a one-way ANOVA followed by Dunnett’s multiple comparisons test relative to the 8-week-old timepoint (I, J), or a Wilcoxon Rank-Sum test (K,L). *P<0.05; **P<0.01; ***P<0.001; ****P<0.0001.

Inactive Hsf1 is typically sequestered in the cytoplasm through physical interactions with chaperones^78–80^. Under proteostasis stress, chaperones dissociate from Hsf1 to bind accumulating unfolded proteins, enabling Hsf1 nuclear translocation and transcriptional induction of chaperones and other proteostasis-stabilizing genes^81,82^. Consistent with this mechanism, immunofluorescence analysis showed robust nuclear Hsf1 in E16.5 HSCs, whereas young adult HSCs displayed predominantly perinuclear staining (Fig. 4F,G). These findings support the view that fetal HSCs operate in a heightened proteostasis demand state that engages Hsf1 even under steady state conditions.

Gene set enrichment (Fig. 4A-E) and gene set variation analysis (Fig. 4H,I) suggested that Hsf1 pathway activity is declining in HSCs at E18.5, is strongly suppressed at P0, and eventually stabilizes at the low homeostatic level characteristic of young adulthood. Notably, Hsf1 transcript abundance remains relatively constant in HSCs across development (Fig. 4J), suggesting that these differences reflect changes in activation state rather than expression.

To determine whether this developmental pattern is conserved in humans, we analyzed published RNA-sequencing datasets from human hematopoietic stem and progenitor cells (HSPCs)^83^. Fetal HSPCs from first and second trimesters exhibited highly significant upregulation of the HSF1 mediated heat shock response signature compared to adult (17–53-year-old) HSPCs (Fig. 4K,L). Together, these findings indicate that fetal HSCs experience heightened proteostasis stress and activate Hsf1 as a conserved adaptive mechanism, linking developmental proteostasis demands to a dedicated stress response program.

### Fetal HSCs transcriptionally rewire the proteostasis network in the absence of Hsf1

To define the role of Hsf1 activation in fetal HSC proteostasis, we conditionally deleted *Hsf1* in the developing hematopoietic system using *Vav1-iCre;Hsf1^fl/fl^* (*Hsf1*^-/-^) mice and examined E16.5 HSPCs. Unexpectedly, *Hsf1* loss did not significantly alter protein synthesis rates or increase unfolded or misfolded protein burden in E16.5 HSCs or progenitor populations (Fig. 5A-C, Supplementary Fig. S3), despite robust activation and expression of related gene sets in wild-type fetal HSCs^73^ (Fig. 4A). However, RNA-sequencing revealed that the absence of overt proteostasis defects belied extensive remodeling of proteostasis pathways. *Hsf1*^-/-^ fetal HSCs showed marked transcriptional induction of protein degradation programs, including both the ubiquitin proteasome system and autophagy, alongside coordinated suppression of ribosome-related gene sets (Fig. 5D-H). These data suggest that fetal HSCs compensate for the loss of *Hsf1* by transcriptionally reprogramming the proteostasis network, activating degradation and restraining anabolic load to maintain proteome integrity. This extensive rewiring suggests that fetal HSCs can transiently preserve proteome stability through Hsf1-independent mechanisms, but this compensation may be intrinsically limited as developmental context shifts.

**Figure 5.**
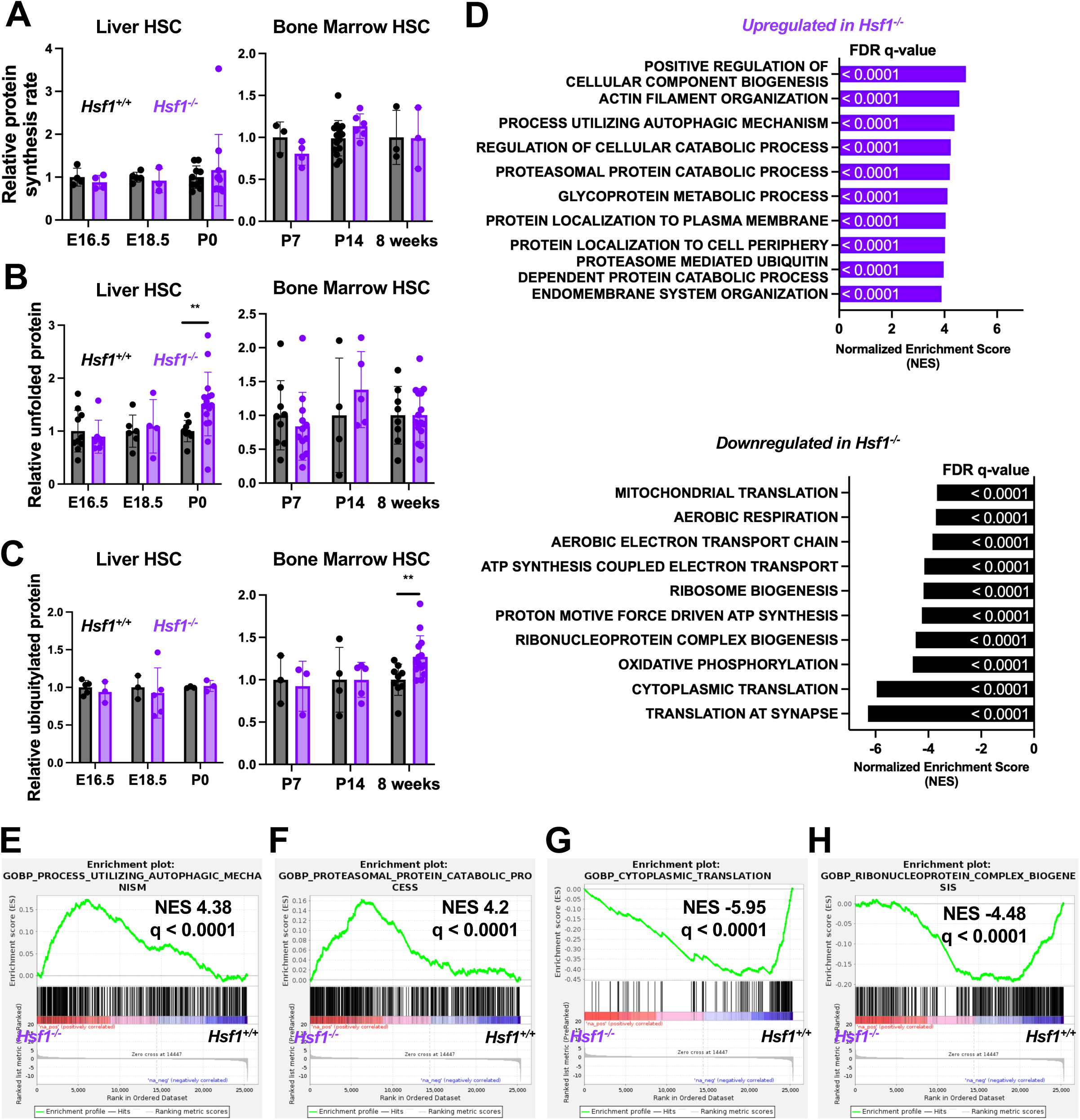
**Fetal HSCs transcriptionally rewire the proteostasis network in the absence of Hsf1** (A) Relative protein synthesis rates in HSCs from *Hsf1^+/+^* and *Hsf1^-/-^* mice at E16.5, E18.5, P0, P7, P14, and 8 weeks. OP-Puro MFI is normalized to *Hsf1^+/+^* HSCs at each respective age (n = 3-14 mice/genotype/age). (B) Relative quantity of unfolded proteins in HSCs in *Hsf1^+/+^* and *Hsf1^-/-^* mice at E16.5, E18.5, P0, P7, P14, and 8 weeks. TMI MFI is normalized to *Hsf1^+/+^* HSCs at each respective age (n = 4-16 mice/genotype/age). (C) Relative quantity of ubiquitylated proteins in HSCs in *Hsf1^+/+^* and *Hsf1^-/-^* mice at E16.5, E18.5, P0, P7, P14, and 8 weeks. MFI is normalized to *Hsf1^+/+^* HSCs at each respective age (n = 3-14 mice/genotype/age). (D) Normalized enrichment scores of the top 10 GOBP upregulated (left) and downregulated (right) pathways in E16.5 *Hsf1^-/-^* as compared to *Hsf1^+/+^* HSCs based on GSEA analysis from RNA sequencing (n=3 mice/genotype). (E-H) Representative enrichment plots of significantly upregulated (E, F) and downregulated (G, H) gene sets related to proteostasis in E16.5 *Hsf1^-/-^* as compared to *Hsf1^+/+^* HSCs. Data represent mean ± standard deviation (A-C). Normalized enrichment scores (NES) and false discovery rates (q) are shown in D-H. Statistical significance was assessed using a Welch’s t test (A-C). **P<0.01.

Because HSC ontogeny involves major, stage-specific transitions in proteostasis, including a sharp reduction in Hsf1 activity at birth (Fig. 4C,H,I), we asked whether this compensatory rewiring during fetal development might render HSCs vulnerable later in maturation. Extending our analyses to later developmental stages revealed that *Hsf1*-deficiency caused a pronounced accumulation of unfolded protein in P0 HSCs (Fig. 5B; P<0.01). This temporal mismatch between early compensatory transcriptional response and later proteostasis instability suggests that fetal adaptations are insufficient to meet the sharply changing proteostasis demands at birth. Thus, while fetal HSCs initially mask the loss of *Hsf1* through extensive transcriptional compensation, this fragile equilibrium transiently collapses as HSCs transition to the neonatal state. However, despite this transient collapse at P0, proteostasis was largely restored at subsequent postnatal stages, indicating that additional maturation dependent mechanisms re-establish homeostasis. Notably, young adult *Hsf1*^-/-^ HSCs exhibited a sustained accumulation of ubiquitylated protein (Fig. 5C), revealing a persistent proteostasis defect that emerges later in life. This finding contrasts with prior reports showing that deleting *Hsf1* in young adulthood does not disrupt proteostasis in young adult HSCs^84^, underscoring the importance of appropriate developmental control of proteostasis and suggesting that fetal loss of Hsf1 induces latent vulnerabilities that manifest later in life.

Together, these findings suggest that Hsf1 is not simply a stress-responsive factor, but a key architect of the fetal proteostasis program. In its absence, fetal HSCs engage a compensatory transcriptional strategy that is ultimately unsustainable, predisposing the HSC proteome to instability at birth and creating latent vulnerabilities that later manifest in young adulthood.

These data therefore establish Hsf1 as a central regulator that ensures continuity of proteostasis across the developmental transition from fetal to neonatal life and safeguards long-term proteome integrity.

### Hsf1 is required for normal fetal hematopoiesis and HSC function

To determine how *Hsf1* loss affects developmental hematopoiesis, we comprehensively analyzed *Hsf1*^-/-^ and littermate control mice across six developmental stages. *Hsf1*-deficiency led to a striking expansion of the stem and progenitor compartments with increased HSC frequency at most timepoints (Fig. 6A-E). This expansion first emerged at E18.5 and peaked at P0, when *Hsf1*^-/-^ mice exhibited 60% more phenotypic HSCs (Fig. 6C; P<0.05). The most primitive MPPs (CD150^-^CD48^-^LSK) similarly increased in frequency at P0 (Fig. 6F), as did the more lineage-biased MPP subsets (Fig. 6G-I). CMPs and MEPs showed more modest expansion, whereas most downstream progenitors and differentiated blood cells remained largely unchanged (Supplementary Fig. S4). Thus, loss of *Hsf1* preferentially perturbs the stem and early progenitor pools during late fetal and neonatal development, consistent with the stage at which proteostasis defects are most pronounced.

**Figure 6.**
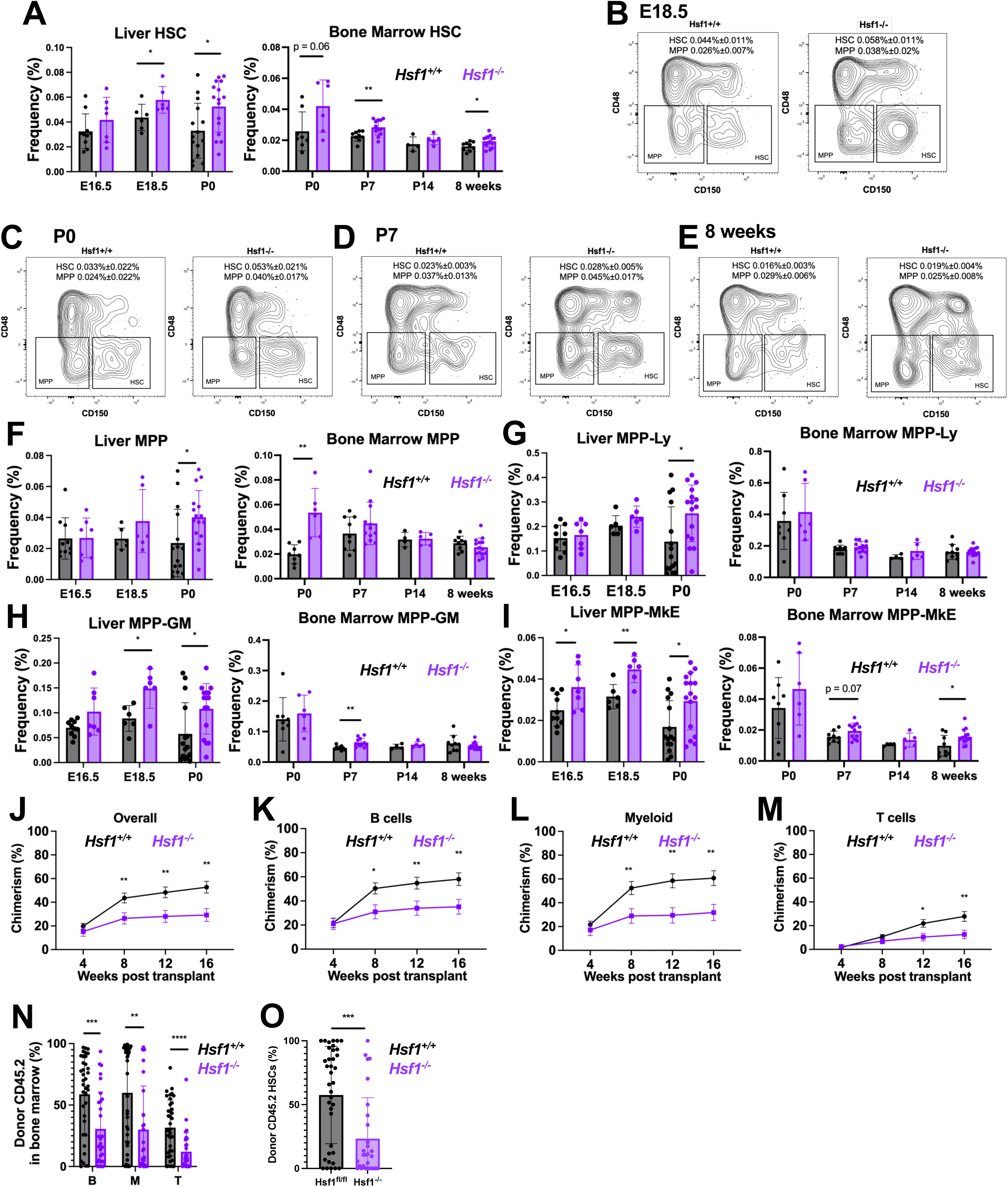
**Hsf1 is required for normal fetal hematopoiesis and HSC function** (A) Frequency of HSCs in *Hsf1^+/+^* and *Hsf1^-/-^* mice at E16.5, E18.5, P0, P7, P14, and 8 weeks (n = 4-16 mice/genotype/age in 1-4 experiments). (B-E) Representative flow cytometry plots and frequency of HSC and MPP populations at E18.5 (B), P0 (C), P7 (D), and 8 weeks (E) in *Hsf1^+/+^* and *Hsf1^-/-^* mice. Plots shown are previously gated on live LSK cells. (F-I) Frequency of MPP (F), MPP-Ly (G), MPP-GM (H), and MPP-MkE (I) populations in *Hsf1^+/+^*and *Hsf1^-/-^* mice at E16.5, E18.5, P0, P7, P14, and 8 weeks (n = 4-16 mice/genotype/age in 1-3 experiments). (J-M) Frequency of donor-derived hematopoietic (J), B (K), myeloid (L), and T (M) cell chimerism when 20 *Hsf1^+/+^* and *Hsf1^-/-^* E16.5 HSCs were transplanted with 3 x 10^5^ bone marrow cells into irradiated mice (n = 38 and 29 recipients of HSCs from *Hsf1^+/+^* and *Hsf1^-/-^* mice, respectively, in 4 total experiments). (N,O) Donor-derived B, myeloid (M), T cells and HSCs (O) in the bone marrow of recipients (from J-M) 16 weeks after transplant. Data represent mean ± standard deviation (A, F-I, N, O) or standard error of the mean (J-M). Differences between groups were assessed by Welch’s t-test. *P<0.05; **P<0.01; ***P<0.001; ****P<0.0001.

We next evaluated how *Hsf1*-deficiency affects the function of fetal HSCs. Transplantation of 20 E16.5 HSCs from *Hsf1*^-/-^ or control mice together with 3x10^5^ wild-type young adult competitor bone marrow cells revealed a pronounced defect in long-term reconstituting activity (Fig. 6J).

Recipients of *Hsf1*^-/-^ fetal HSCs showed significantly reduced donor chimerism in peripheral blood and bone marrow B-, T-, and myeloid lineages (Fig. 6K-N), accompanied by a significant decrease in donor-derived HSCs in the recipient bone marrow (Fig. 6O). Notably, this developmental requirement for Hsf1 is stage-specific, as our prior work and studies from others demonstrated that Hsf1 is fully dispensable for young adult HSC function^77,84,85^. These defects in *Hsf1*^-/-^ fetal HSCs likely reflect the cumulative impact of operating in a compensated proteostasis state during development, rather than an acute proteostasis failure at the time of transplantation.

Coupled with the transcriptional compensation and neonatal proteostasis collapse described above, these findings reveal that Hsf1 is essential for aligning proteostasis capacity with the functional demands of fetal HSCs. In fetal HSCs, Hsf1 supports appropriate proteostasis architecture, and its absence triggers compensatory expansion and functional impairment.

Together these data position Hsf1 as a central and developmentally tuned determinant of both proteome integrity and functional competence during HSC maturation.

## DISCUSSION

Our study identifies a developmental shift in proteostasis control that distinguishes fetal, neonatal, and young adult HSCs. We show that fetal HSCs operate under an intrinsically elevated burden of protein synthesis and unfolded and misfolded proteins, accompanied by robust activation of Hsf1. This contrasts sharply with young adult HSCs that maintain a more quiescent proteostasis state marked by very low protein synthesis rates coupled with heightened protein quality^2,3^. We found that protein synthesis rates decline progressively across ontogeny, and this developmental gradient highlights a dynamic and age-dependent rewiring of the proteostasis landscape. Together, these findings suggest that early hematopoietic development tolerates a unique proteostasis configuration that supports rapid expansion but increases reliance on dedicated stress-mitigating pathways.

Our genetic studies reveal that Hsf1 is not simply an inducible stress-response factor in HSCs^77^, but a developmental architect that establishes the fetal HSC proteostasis program. The dispensability of Hsf1 in young adult HSCs^77,84^ highlights an ontogeny-specific dependency that was previously unappreciated. Strikingly, fetal HSCs initially compensate for *Hsf1* loss, as E16.5 *Hsf1*^-/-^ HSCs maintain normal protein synthesis rates and do not accumulate unfolded protein.

This “compensation without stress” phenotype is accompanied by transcriptional rewiring of the proteostasis network, including upregulation of proteasome and autophagy pathways. However, the neonatal transition marks a vulnerability window in which the essentiality of Hsf1 is revealed, leading to accumulation of unfolded protein. Interestingly, neonatal HSCs that normally suppress Hsf1 activity, exhibit heightened proteasome capacity and dependency, suggesting that elevated proteasomal flux is a critical buffer when protein synthesis rates are high and Hsf1 activity is suppressed. This reveals remarkable plasticity in the developmental proteostasis network.

Disruption of proteostasis integrity also alters the architecture of the hematopoietic stem and progenitor compartment. Fetal and neonatal *Hsf1*^-/-^ mice show selective expansion of HSCs and early MPPs, which may reflect a compensatory attempt to preserve hematopoietic output as proteostasis destabilizes. The sharp decline in long-term multilineage reconstituting activity in fetal *Hsf1*^-/-^ HSCs underscores how tightly HSC pool size and lifelong functionality are coupled to proteostasis control. More broadly, the differential requirement for Hsf1 across developmental stages highlights that proteostasis regulators are not uniformly essential across stem cell states, raising the possibility that other canonical stress response pathways may also have developmentally restricted roles.

These results also provide a framework for understanding why early life HSCs may be particularly vulnerable to perturbations in proteostasis. Proteostasis disruption is a unifying mechanism across inherited bone marrow failure syndromes with diverse mutations^86^, and extends beyond classical ribosomopathies that impair ribosomal biogenesis^87–95^ and translation initiation^89,96–100^ to encompass disorders with mutations that impair protein processing and trafficking^101–111^, RNA processing^93,112^, translation^113–115^, and DNA damage repair^55,116–121^. Our findings suggest that the high protein biosynthetic load and proteostasis demands characteristic of fetal and neonatal HSCs may exacerbate the functional consequences of such mutations, rendering early-life HSCs disproportionately sensitive to even modest impairments in proteostasis machinery. This developmental vulnerability aligns with the early childhood onset of marrow failure and heightened and progressive susceptibility to clonal hematopoiesis and leukemia predisposition in select inherited bone marrow failure syndromes and related germline disorders. These principles may also help explain spontaneous remissions that are occasionally observed in some of these disorders, as developmental changes in proteostasis load and regulation could rebalance destabilized stem cell states. Defining ontogenic proteostasis dependencies could reveal opportunities to develop therapies that stabilize the stressed developing proteome or selectively target aberrant proteostasis states to improve outcomes in pediatric bone marrow failure and preleukemic conditions.

The developmental Hsf1 program uncovered here also resonates with our work in aging HSCs, where Hsf1 activation contributes to maladaptive stress remodeling^84^. Understanding how Hsf1 activation is precisely regulated or dysregulated during these developmental windows may offer insight into why some pediatric leukemias originate from fetal or neonatal HSCs, and how Hsf1 activity could cooperate with oncogenic lesions during these windows of heightened proteostasis strain.

Several limitations of this study should be acknowledged. First, while Hsf1 clearly serves as a central proteostasis node, we have not yet delineated the full network of parallel or compensatory pathways, including proteasome regulation, autophagy, and the unfolded protein response, and the mechanisms that govern their developmental deployment remain unidentified. Second, our analyses largely capture population-level proteostasis states and may obscure potentially meaningful heterogeneity within fetal and neonatal HSCs. Third, we evaluated select stages of fetal development. Evaluating earlier fetal ages may reveal a more complete trajectory of early developmental changes in proteostasis pathways, though cell numbers may be technically limiting. Finally, although our findings highlight strong conceptual links to inherited bone marrow failure syndromes, we did not directly test disease-associated mutations in the fetal or neonatal context. These limitations identify concrete and mechanistically tractable areas for future investigation.

By defining Hsf1 as a developmental regulator of the fetal HSC proteostasis network, revealing transient plasticity that masks early deficits and identifying birth as a critical vulnerability window, this work positions proteostasis integrity as a foundational determinant of lifelong HSC competence. These insights bridge fundamental stem cell biology with clinical questions in pediatric bone marrow failure and leukemia predisposition, laying the groundwork for future mechanistic and translational studies.

## METHODS

### Mice

Female and male C57BL6/J mice were analyzed at E16.5, E18.5, P0 (birth), P7, P14, and 7-10 weeks old (young adult). CAG-RFP-EGFP-LC3 mice^122^ were maintained by breeding CAG-RFP-EGFP-LC3/+ mice with C57BL6/J mice. Female *Vav-iCre* mice^123^ were obtained from The Jackson Laboratory (Strain #008610) and crossed to male *Hsf1^fl/fl^* mice^124^. Female *Vav-iCre;Hsf1^fl/fl^* were bred with male *Hsf1^fl/fl^* mice in timed matings to generate experimental litters.

All mice were housed in the vivarium at the University of California San Diego Moores Cancer Center or Sanford Consortium for Regenerative Medicine in specific pathogen-free conditions. All protocols were approved by the University of California San Diego Institutional Animal Care and Use Committee.

### Cell Isolation

Livers were dissected from E16.5, E18.5, and P0 mice and were crushed between frosted slides and passed through a 40 μm cell strainer prior to staining for each assay. Bone marrow was isolated from long bones of P0, P7, P14, and young adult mice. When additional cells were required, bone marrow was also obtained from crushed and filtered spine and hip bones. Cells were processed in Ca^2+^- and Mg^2+^-free Hank’s buffered salt solution (HBSS; Corning 21-022-CV) supplemented with 2% (v/v) heat-inactivated bovine serum (Gibco 26170043). Cell number and viability were assessed with a hemocytometer or TC20 Automated Cell Counter (BioRad) based on trypan blue exclusion.

### Flow cytometry and cell sorting

For flow cytometric analysis and isolation of specific hematopoietic populations, cells were incubated with combinations of antibodies to the following cell-surface markers, conjugated to FITC, PE, PerCP-Cy5.5, APC, APC-eFluor 780, PE-Cy7, Alexa Fluor 700, or biotin (antibody clones are given in brackets in the following list): CD3 (17A2), CD4 (GK1.5), CD5 (53-7.3), CD8α (53-6.7), CD11b (M1/70), CD16/32 (FcγRII/III; 93), CD34 (RAM34), CD41 (MWReg30), CD43 (R2/60), CD45.1 (A20), CD45.2 (104), CD45R (B220; RA3-6B2), CD48 (HM48-1), CD71 (R17217), CD117 (c-Kit; 2B8), CD127 (IL7Rα; A7R34), CD135 (A2F10), CD150 (TC15-12F12.2), Ter119 (TER-119), Sca-1 (D7, E13-161.7), Gr-1 (RB6-8C5), and IgM (II/41). Biotinylated antibodies were visualized by incubation with PE-Cy7 conjugated streptavidin (BioLegend 405206). All reagents were acquired from eBiosciences (Thermo Fisher Scientific) or BioLegend. All incubations were for 30-60 minutes on ice.

For isolation and analysis of mouse HSC and MPP populations, Lineage markers included CD3, CD5, CD8α, B220, Gr-1, and Ter119. HSCs were defined as CD150^+^CD48^−^Lineage^−^Sca1^+^cKit^+59^, MPP as CD150^−^CD48^−^Lineage^−^Sca1^+^cKit^+65,125^, MPP^G/M^ as CD150^−^CD48^+^Lineage^−^Sca1^+^cKit^+65,126^, MPP^Mk/E^ as CD150^+^CD48^+^Lineage^−^Sca1^+^cKit^+65,126^ and MPP^Ly^ as CD135^+^Lineage^−^Sca1^+^cKit^+65,126^. For analysis of mouse CMPs, GMPs, and MEPs, Lineage markers included CD3, CD4, CD8α, CD11b, B220, Gr-1, and Ter119. CMPs were defined as CD34^+^CD16/32^low^CD127^−^Lineage^−^Sca1^−^cKit^+66^, GMPs as CD34^+^CD16/32^high^CD127^−^Lineage^−^Sca1^−^cKit^+66^ and MEPs as CD34^−^CD16/32^low^CD127^−^Lineage^−^Sca1^−^cKit^+66^. For lymphoid lineages, IgM^+^ B cells were defined as IgM^+^B220^+127^, Pre-B cells as IgM^−^CD43^−^B220^+127^, Pro-B cells as IgM^−^CD43^+^B220^+127^ and T cells as CD4^+^ or CD8^+^. For myeloid lineages, erythrocytes were defined as Ter119^+^, myeloid cells (GM) as Gr1^+^CD11b^+^ and megakaryocytes (Mk) as CD41^+^.

HSCs were pre-enriched by selecting c-Kit^+^ cells using paramagnetic microbeads and an autoMACS magnetic separator (Miltenyi Biotec) before sorting. Non-viable cells were excluded from sorts and analyses using 1 μg/mL DAPI.

Cell sorting was performed on a BD FACSAriaII or BD FACSAria Fusion. Sorted fractions were double sorted (yield then purity precision) or sorted with single cell precision to ensure high purity. Data acquisition for analysis was performed using BD FACSymphony A1. Data were analyzed using FlowJo (BD) software.

### Protein synthesis

Protein synthesis was quantified using O-propargyl-puromycin (OP-Puro). OP-Puro (50 mg/kg; Medchem Source) was administered by intraperitoneal injection into neonatal, young adult mice, or pregnant females at the indicated gestational stages. One hour later, liver or bone marrow cells were harvested, stained with antibodies, and OP-Puro incorporated peptides were detected via click-chemistry labeling and quantified by flow cytometry as previously described^2,57^. Fluorescence intensity of each HSPC population was normalized to DAPI^+^ cells from the same sample, and relative protein synthesis was calculated compared with young adults or controls.

### Unfolded protein

Liver or bone marrow cells were stained for cell surface markers as described above. After staining, cells were washed twice in Ca^2+^- and Mg^2+^-free PBS. Tetraphenylethene maleimide (TMI^67^; stock 2 mM in DMSO) was diluted in PBS (50 μM final concentration), added to each sample and samples were incubated at 37°C for 45 minutes. Samples were washed twice in PBS and analyzed by flow cytometry. DAPI was omitted from these samples because of spectral overlap. Relative TMI mean fluorescence intensity of each age normalized to unfractionated cells compared to 8-week-old mice or control mice was calculated.

### Ubiquitylated protein

For analysis of ubiquitylated protein following cell surface staining, cells were fixed in 0.5 ml of 1% paraformaldehyde (Affymetrix) in PBS for 10-15 min on ice. Cells were washed in PBS, then permeabilized in 200 ml PBS supplemented with 3% (v/v) fetal bovine serum (Thermo Fisher Scientific) and 0.1% (m/v) saponin (Sigma) for 5 min at room temperature (20–25°C). Cells were then incubated with anti-ubiquitylated protein antibody (clone FK2, Millipore) at a concentration of 1:500 for 30 min at room temperature. This was followed by incubation with anti-mouse Alexa Fluor 488 (Thermo Fisher Scientific) at a concentration of 1:500 for 30 min at room temperature. Cells were washed twice in PBS supplemented with 3% fetal bovine serum and 0.1% saponin, then resuspended in PBS supplemented with DAPI (4 mg/ml final concentration) and analyzed by flow cytometry. Relative fluorescence (mean fluorescence intensity) of each age normalized to DAPI^+^ cells compared to 8-week-old mice or control mice was calculated.

### Proteasome activity

The Proteasome-Glo assay was used to measure proteasome activity in purified HSPCs as previously described^3,4,77^. The Proteasome-Glo Chymotrypsin-Like Cell-Based Assay reagent (Promega) was applied to pooled groups of multiples of 5 x 10^3^ LSK, CMP, GMP, and MEP cells in solid opaque 96-well plates. A Tecan Spark plate reader was used to measure luminescence of each group of cells.

### Proteasome inhibition

Mice received an intraperitoneal injection of 10 mg/kg bortezomib (Cell Signaling) diluted in 10% (v/v) DMSO. The spine and long bones were harvested 2 hours later for analysis.

### Transplantation assays

Adult recipient mice were irradiated with two doses of 500 cGy 4 hours apart using an X-Rad320 X-ray irradiator (Precision X-Ray) to provide a total dose of 1000 cGy. 20 donor HSCs from E16.5 *Vav-iCre;Hsf1^fl/fl^* and *Hsf1^fl/fl^* mice were injected into the retro-orbital venous sinus of anesthetized, irradiated CD45.1 recipient mice with 3x10^5^ recipient-type young adult bone marrow cells. Recipient mice were bled every 4 weeks for 16 weeks after transplantation. Red blood cells were lysed with ammonium chloride potassium buffer. The remaining cells were stained with antibodies against CD45.2, CD45.1, CD45R (B220), CD11b, CD3, and Gr-1 to assess donor-cell engraftment (chimerism). After sixteen weeks post-transplantation, bone marrow of primary recipients was analyzed for HSC engraftment and multilineage chimerism. Long-term multilineage reconstitution was achieved when the recipient had ≥ 0.5% donor-derived peripheral B, T, and myeloid cells.

### RNA-sequencing

For RNA-sequencing analysis, total RNA was extracted from up to 3,104 HSCs per embryo using the RNeasy Plus Micro Kit (Qiagen 74034). Illumina mRNA libraries were prepared using the SMART-seq protocol^128^ (Takara Bio). 2-2.6μL of total RNA was used in the SMART-seq protocol (100-400pg). 10 cycles of PCR were performed for the cDNA preamplification step, and 18 cycles were performed for the tagmentation library preparation.

The resulting libraries were double-sided size selected (0.6-1x), pooled and deep sequenced in one lane on the Illumina NovaSeq X Plus using paired-end reads with both forward and reverse read lengths of 75 nucleotides (80-107M reads per condition). The reads that passed Illumina filters were filtered for reads aligning to tRNA, rRNA, adapter sequences, and spike-in controls. The reads were then aligned to GRCm38 reference genome using STAR (v2.6.1c)^129^. DUST scores were calculated with PRINSEQ Lite (v 0.20.3)^130^ and low complexity reads (DUST > 4) were removed from the BAM files. The alignments results were parsed via SAMtools^131^ to generate SAM files. Read counts to each genomic feature were obtained with htseq-count program (v0.7.1)^132^ using the “union” option. After removing absent features (zero counts in all samples), the raw counts were then imported to Bioconductor package DESeq2 (v1.24.0)^133^ to identify differentially expressed genes among samples. P-values for differential expression were calculated using the Wald test for differences between the base means of two conditions. These P-values were then adjusted for multiple test correction using Benjamini Hochberg algorithm.

We considered genes differentially expressed between two groups of samples when the DESeq2 analysis resulted in an adjusted P-value of <0.05 and the difference in gene expression was at least 1.5-fold.

GSEA^134^ was done on GSE128759 and E16.5 *Vav-iCre;Hsf1^fl/fl^* HSCs using the “GseaPreranked” method with “classic” scoring scheme. MSigDB gene sets^135,136^ were downloaded for mouse Gene Ontology^137^ analysis. Rank files for each DESeq2 comparison were generated by assigning a rank of negative log10(pValue) to genes with log2FoldChange greater than zero and a rank of positive log10(pValue) to genes with a log2FoldChange less than zero.

To assess the heat shock response activity in HSCs across various developmental stages, GSVA^138^ was performed on the log2CPM data from dataset GSE128759 and MSigDB gene sets using the GSEApy package^139^. The resulting GSVA enrichment scores were visualized using heatmaps and violin plots.

### Hsf1 immunostaining and microscope image acquisition

For analysis of Hsf1 activation, freshly isolated HSCs were sorted and plated on VistaVision HistoBond Microscope Slides (VWR 16004-406) with RetroNectin (Clontech Labs/Takara Bio T100A). RetroNectin was diluted to 12.5μg/mL in DPBS (Corning 21-030-CV) and 5μL was placed for each sample on top of microscope slides. To allow RetroNectin to bind, slides were incubated at 37°C for at least 30 minutes in a humid environment. Excess RetroNectin was removed before plating cells on top of the RetroNectin droplet to allow cells to bind to the microscope slide. RetroNectin slides with cells were incubated at 37°C for 15 minutes. Cells were then directly fixed in 4% paraformaldehyde in PBS (Electron Microscopy Sciences 157-4) for 15 minutes at room temperature. Cells were washed three times with PBS and permeabilized with PBS supplemented with 0.1% Triton X-100 (VWR 80503-490), 0.03% Tween20 (Sigma P9416) and 1% BSA (Sigma A9647) for 20 minutes at room temperature. Cells were washed three times with PBS supplemented with 0.03% Tween20. Cells were blocked with PBS supplemented with 1% BSA and 0.03% Tween20 for 60 minutes at room temperature. Samples were incubated with anti-Hsf1 antibody (Cell Signaling, Rabbit polyclonal 4356) at 1:500 dilution in blocking solution overnight at 4°C. Cells were washed three times with PBS supplemented with 0.03% Tween20 and incubated with Alexa Fluor 488 conjugated donkey anti-Rabbit secondary antibody (Thermo Fisher A-21206) at 1:500 dilution in blocking solution with 2μg/mL DAPI for 60 minutes at room temperature. Cells were washed three times with PBS and mounted in ProLong Diamond Antifade Mountant (Molecular Probes P36961) with No. 1.5 Gold Seal Cover Glass (Electron Microscopy Sciences 6379110). All processes were done on the RetroNectin microscope slides with a hydrophobic barrier (Newcomer Supply PAP Pen Liquid Blocker 6506).

Images were acquired using a DMi8 microscope (Leica) attached to a spinning disk Dragonfly 200 High Speed Confocal Microscope System (Andor/Oxford Instruments) with the Fusion software (Andor). All images were acquired using a 63X oil-immersion objective with Zyla cameras and excited using 405nm (DAPI) and 488nm (Alexa Fluor 488) lasers. Z-stack images were taken with the same confocal settings for Alexa Fluor 488 with minor adjustments for DAPI between samples. Raw images were processed with FIJI (ImageJ) and imported as a Hyperstack. Max Intensity ZProjection was generated for each Hyperstack. Blue (DAPI) and green (anti-Hsf1/Alexa Fluor 488) channels were merged, and Alexa Fluor 488 intensity was quantified based on co-localization of DAPI and Hsf1. RawIntDen/Area was used for quantification and at least 100 cells were quantified per sample if possible. The same Brightness/Contrast settings were used across samples within an experiment for Alexa Fluor 488 with minor adjustments for DAPI between samples.

### Statistical analysis

Data are represented by mean ± standard deviation (SD), except for transplantation data which are represented as mean ± standard error of the mean (SEM). To test statistical significance between two samples, two-tailed Welch’s t-tests were used. When multiple samples were compared to one another, statistical significance was assessed using repeated-measures one-way ANOVA followed by Dunnett’s test for multiple comparisons.

Statistical tests were performed using GraphPad Prism software.

## ACKNOWLEDGMENTS

Work in the Signer Laboratory is supported by the NIH/NIDDK (R01DK116951), NIH/NHLBI (R01HL165172, R01HL180754), NIH/NCI (U01CA267031), NIH/NIA (R01AG088725), the American Cancer Society (CSCC-RSG-23-994830-01-CSCC), the Sanford Stem Cell Institute, the UC San Diego Moores Cancer Center which is supported by the NCI Cancer Center Support Grant (P30CA023100), and an anonymous private family foundation. HY and RAJS were supported by a Curebound Discovery Grant. HY is supported by NIH/NICHD (5T32HD087978), the Pediatric Transplantation and Cellular Therapy Consortium New Investigator Award funded by the PBMTF, Hyundai Hope on Wheels, Alex’s Lemonade Stand Foundation (1416965), and the American Association for Cancer Research (25-40-64-YU). The LJI Flow Cytometry Core, LJI Sequencing Core Facility, UCSD IGM Genomics Center, and UCSD Moores Cancer Center flow cytometry facility are supported by the NIH Shared Instrumentation Grant Program (S10 RR027366, S10 OD025052, S10 OD026929, S10 OD032316). The Sanford Consortium for Regenerative Medicine (SCRM) Flow Core is part of the UCSD Human Embryonic Stem Cell Core Facility and is supported by a CIRM Major Facilities grant (FA1-00607). We would like to thank the LJI Bioinformatics core and the Stem Cell Genomics Core at SCRM for providing bioinformatics, sequencing, and microscopy services, E. Bennett and J. Magee for advice, Y. Hong for TMI, and E. Christians for *Hsf1*^fl^ mice.

## AUTHOR CONTRIBUTIONS

H.Y. and R.A.J.S. conceived the project, designed experiments, and wrote the manuscript. H.Y. performed and analyzed most experiments. Y.K. performed some computational analyses. K.C. provided technical support. A.Z.L. performed microscopy experiments. M.S. supported mouse breeding and laboratory operations.

## AUTHOR INFORMATION

The authors declare no competing interests.

## REFERENCES

1 Chua, B. A. & Signer, R. A. J. Hematopoietic stem cell regulation by the proteostasis network. Current Opinion in Hematology 27, 254–263 (2020). 10.1097/MOH.0000000000000591

2 Signer, R. A. J., Magee, J. A., Salic, A. & Morrison, S. J. Haematopoietic stem cells require a highly regulated protein synthesis rate. Nature 508, 49–54 (2014). 10.1038/nature13035

3 Hidalgo San Jose, L., et al. Modest Declines in Proteome Quality Impair Hematopoietic Stem Cell Self-Renewal. Cell Reports 30, 69–80.e66 (2020). 10.1016/j.celrep.2019.12.003

4 Chua, B. A. et al. Hematopoietic stem cells preferentially traffic misfolded proteins to aggresomes and depend on aggrephagy to maintain protein homeostasis. Cell Stem Cell 30, 460–472.e466 (2023). 10.1016/j.stem.2023.02.010

5 Van Galen, P. et al. The unfolded protein response governs integrity of the haematopoietic stem-cell pool during stress. Nature 510, 268–272 (2014). 10.1038/nature13228

6 Herrejon Chavez, F., et al. RNA binding protein SYNCRIP maintains proteostasis and self-renewal of hematopoietic stem and progenitor cells. Nature Communications 14 (2023). 10.1038/s41467-023-38001-x

7 Chapple, R. H. et al. ERα promotes murine hematopoietic regeneration through the ire1α-mediated unfolded protein response. eLife 7, 1–25 (2018). 10.7554/eLife.31159

8 Xie, S. Z. et al. Sphingolipid Modulation Activates Proteostasis Programs to Govern Human Hematopoietic Stem Cell Self-Renewal. Cell Stem Cell 25, 639–653.e637 (2019). 10.1016/j.stem.2019.09.008

9 Signer, R. A. J. et al. The rate of protein synthesis in hematopoietic stem cells is limited partly by 4E-BPs. Genes and Development 30, 1698–1703 (2016). 10.1101/gad.282756.116

10 Li, C. et al. Amino acid catabolism regulates hematopoietic stem cell proteostasis via a GCN2-eIF2α axis. Cell Stem Cell 29, 1119–1134.e1117 (2022). 10.1016/j.stem.2022.06.004

11 Moran-Crusio, K., Reavie, L. B. & Aifantis, I. Regulation of hematopoietic stem cell fate by the ubiquitin proteasome system. Trends in Immunology 33, 357–363 (2012). 10.1016/j.it.2012.01.009

12 Thompson, B. J. et al. Control of hematopoietic stem cell quiescence by the E3 ubiquitin ligase Fbw7. Journal of Experimental Medicine 205, 1395–1408 (2008). 10.1084/jem.20080277

13 King, B. et al. The ubiquitin ligase Huwe1 regulates the maintenance and lymphoid commitment of hematopoietic stem cells. Nature Immunology 17, 1312–1321 (2016). 10.1038/ni.3559

14 Reavie, L. et al. Regulation of hematopoietic stem cell differentiation by a single ubiquitin ligase-substrate complex. Nature Immunology 11, 207–215 (2010). 10.1038/ni.1839

15 Ho, T. T. et al. Autophagy maintains the metabolism and function of young and old stem cells. Nature 543, 205–210 (2017). 10.1038/nature21388

16 van Galen, P. et al. Integrated Stress Response Activity Marks Stem Cells in Normal Hematopoiesis and Leukemia. Cell Reports 25, 1109–1117.e1105 (2018). 10.1016/j.celrep.2018.10.021

17 Bowie, M. B. et al. Hematopoietic stem cells proliferate until after birth and show a reversible phase-specific engraftment defect. Journal of Clinical Investigation 116, 2808–2816 (2006). 10.1172/JCI28310

18 Bowie, M. B. et al. Identification of a new intrinsically timed developmental checkpoint that reprograms key hematopoietic stem cell properties. Proceedings of the National Academy of Sciences of the United States of America 104, 5878–5882 (2007). 10.1073/pnas.0700460104

19 Nygren, J. M., Bryder, D. & Jacobsen, S. E. Prolonged cell cycle transit is a defining and developmentally conserved hemopoietic stem cell property. J Immunol 177, 201–208 (2006). 10.4049/jimmunol.177.1.201

20 Morrison, S. J., Hemmati, H. D., Wandycz, A. M. & Weissman, I. L. The purification and characterization of fetal liver hematopoietic stem cells. Proceedings of the National Academy of Sciences of the United States of America 92, 10302–10306 (1995). 10.1073/pnas.92.22.10302

21 Maillard, L. et al. CD117^hi^ expression identifies a human fetal hematopoietic stem cell population with high proliferation and self-renewal potential. Haematologica 105, e43–e47 (2020). 10.3324/haematol.2018.207811

22 Pietras, E. M., Warr, M. R. & Passegué, E. Cell cycle regulation in hematopoietic stem cells. J Cell Biol 195, 709–720 (2011). 10.1083/jcb.201102131

23 Manesia, J. K. et al. Highly proliferative primitive fetal liver hematopoietic stem cells are fueled by oxidative metabolic pathways. Stem Cell Research 15, 715–721 (2015). 10.1016/j.scr.2015.11.001

24 Ansó, E. et al. The mitochondrial respiratory chain is essential for haematopoietic stem cell function. Nat Cell Biol 19, 614–625 (2017). 10.1038/ncb3529

25 Nakamura-Ishizu, A., Ito, K. & Suda, T. Hematopoietic Stem Cell Metabolism during Development and Aging. Developmental Cell 54, 239–255 (2020). 10.1016/j.devcel.2020.06.029

26 Ito, K. et al. Self-renewal of a purified Tie2+ hematopoietic stem cell population relies on mitochondrial clearance. Science 354, 1156–1160 (2016). 10.1126/science.aaf5530

27 Dong, S. et al. Chaperone-mediated autophagy sustains haematopoietic stem-cell function. Nature 591, 117–123 (2021). 10.1038/s41586-020-03129-z

28 Grant Rowe, R., et al. Developmental regulation of myeloerythroid progenitor function by the Lin28b-let-7-Hmga2 axis. Journal of Experimental Medicine 213, 1497–1512 (2016). 10.1084/jem.20151912

29 Mold, J. E. et al. Fetal and Adult Hematopoietic Stem Cells Give Rise to Distinct T Cell Lineages in Humans. Science 330, 1695–1700 (2010).

30 Benz, C. et al. Hematopoietic stem cell subtypes expand differentially during development and display distinct lymphopoietic programs. Cell Stem Cell 10, 273–283 (2012). 10.1016/j.stem.2012.02.007

31 Beaudin, A. E. et al. A Transient Developmental Hematopoietic Stem Cell Gives Rise to Innate-like B and T Cells. Cell Stem Cell 19, 768–783 (2016). 10.1016/j.stem.2016.08.013

32 Ganuza, M. et al. Murine foetal liver supports limited detectable expansion of life-long haematopoietic progenitors. Nat Cell Biol 24, 1475–1486 (2022). 10.1038/s41556-022-00999-5

33 Ulloa, B. A. et al. Definitive hematopoietic stem cells minimally contribute to embryonic hematopoiesis. Cell Rep 36, 109703 (2021). 10.1016/j.celrep.2021.109703

34 Yokomizo, T. et al. Independent origins of fetal liver haematopoietic stem and progenitor cells. Nature 609, 779–784 (2022). 10.1038/s41586-022-05203-0

35 Notta, F. et al. Distinct routes of lineage development reshape the human blood hierarchy across ontogeny. Science 351, aab2116 (2016). 10.1126/science.aab2116

36 Kristiansen, T. A. et al. Developmental cues license megakaryocyte priming in murine hematopoietic stem cells. Blood Adv 6, 6228–6241 (2022). 10.1182/bloodadvances.2021006861

37 Copley, M. R. et al. The Lin28b-let-7-Hmga2 axis determines the higher self-renewal potential of fetal haematopoietic stem cells. Nature Cell Biology 15, 916–925 (2013). 10.1038/ncb2783

38 Sigurdsson, V. et al. Bile Acids Protect Expanding Hematopoietic Stem Cells from Unfolded Protein Stress in Fetal Liver. Cell Stem Cell 18, 522–532 (2016). 10.1016/j.stem.2016.01.002

39 Warr, M. R. et al. FOXO3A directs a protective autophagy program in haematopoietic stem cells. Nature 494, 323–327 (2013). 10.1038/nature11895

40 Baldridge, M. T., King, K. Y., Boles, N. C., Weksberg, D. C. & Goodell, M. A. Quiescent haematopoietic stem cells are activated by IFN-gamma in response to chronic infection. Nature 465, 793–797 (2010). 10.1038/nature09135

41 Essers, M. A. et al. IFNalpha activates dormant haematopoietic stem cells in vivo. Nature 458, 904–908 (2009). 10.1038/nature07815

42 Pietras, E. M. et al. Re-entry into quiescence protects hematopoietic stem cells from the killing effect of chronic exposure to type I interferons. The Journal of experimental medicine 211, 245–262 (2014). 10.1084/jem.20131043

43 Chavez, J. S. et al. PU.1 enforces quiescence and limits hematopoietic stem cell expansion during inflammatory stress. J Exp Med 218 (2021). 10.1084/jem.20201169

44 Magee, J. A. & Signer, R. A. J. Developmental Stage-Specific Changes in Protein Synthesis Differentially Sensitize Hematopoietic Stem Cells and Erythroid Progenitors to Impaired Ribosome Biogenesis. Stem Cell Reports 16, 20–28 (2021). 10.1016/j.stemcr.2020.11.017

45 McKinney-Freeman, S. et al. The transcriptional landscape of hematopoietic stem cell ontogeny. Cell Stem Cell 11, 701–714 (2012). 10.1016/j.stem.2012.07.018

46 Okeyo-Owuor, T. et al. The efficiency of murine MLL-ENL-driven leukemia initiation changes with age and peaks during neonatal development. Blood Advances 3, 2388–2399 (2019). 10.1182/bloodadvances.2019000554

47 Mendoza-Castrejon, J. & Magee, J. A. Layered immunity and layered leukemogenicity: Developmentally restricted mechanisms of pediatric leukemia initiation. Immunological Reviews 315, 197–215 (2023). 10.1111/imr.13180

48 Porter, S. N. et al. Fetal and neonatal hematopoietic progenitors are functionally and transcriptionally resistant to. Elife 5 (2016). 10.7554/eLife.18882

49 Lopez, C. K. et al. Ontogenic Changes in Hematopoietic Hierarchy Determine Pediatric Specificity and Disease Phenotype in Fusion Oncogene-Driven Myeloid Leukemia. Cancer Discov 9, 1736–1753 (2019). 10.1158/2159-8290.CD-18-1463

50 Horton, S. J. et al. MLL-AF9-mediated immortalization of human hematopoietic cells along different lineages changes during ontogeny. Leukemia 27, 1116–1126 (2013). 10.1038/leu.2012.343

51 Chen, W., O’Sullivan, M. G., Hudson, W. & Kersey, J. Modeling human infant MLL leukemia in mice: leukemia from fetal liver differs from that originating in postnatal marrow. Blood 117, 3474–3475 (2011). 10.1182/blood-2010-11-317529

52 Mendoza-Castrejon, J. et al. Fetal context conveys heritable protection against MLL-rearranged AML that depends on MLL3. Blood (2025). 10.1182/blood.2025029686

53 Li, Y. et al. LIN28B promotes differentiation of fully transformed AML cells but is dispensable for fetal leukemia suppression. Leukemia 38, 648–651 (2024). 10.1038/s41375-024-02167-0

54 Eldeeb, M. et al. A fetal tumor suppressor axis abrogates MLL-fusion-driven acute myeloid leukemia. Cell Rep 42, 112099 (2023). 10.1016/j.celrep.2023.112099

55 Kovuru, N. et al. Deregulated protein homeostasis constrains fetal hematopoietic stem cell pool expansion in Fanconi anemia. Nature Communications 15, 1–15 (2024). 10.1038/s41467-024-46159-1

56 Wang, K. et al. Ontogeny Dictates Oncogenic Potential, Lineage Hierarchy, and Therapy Response in Pediatric Leukemia. *Cancer Discov*, OF1-OF30 (2025). 10.1158/2159-8290.CD-25-0556

57 Hidalgo San Jose, L. & Signer, R. A. J. Cell-type-specific quantification of protein synthesis in vivo. Nature Protocols 14, 441–460 (2019). 10.1038/s41596-018-0100-z

58 Liu, J., Xu, Y., Stoleru, D. & Salic, A. Imaging protein synthesis in cells and tissues with an alkyne analog of puromycin. Proceedings of the National Academy of Sciences of the United States of America 109, 413–418 (2012). 10.1073/pnas.1111561108

59 Kiel, M. J. et al. SLAM family receptors distinguish hematopoietic stem and progenitor cells and reveal endothelial niches for stem cells. Cell 121, 1109–1121 (2005). 10.1016/j.cell.2005.05.026

60 Kikuchi, K. & Kondo, M. Developmental switch of mouse hematopoietic stem cells from fetal to adult type occurs in bone marrow after birth. Proceedings of the National Academy of Sciences of the United States of America 103, 17852–17857 (2006). 10.1073/pnas.0603368103

61 Hall, T. D. et al. Murine fetal bone marrow does not support functional hematopoietic stem and progenitor cells until birth. Nature Communications 13 (2022). 10.1038/s41467-022-33092-4

62 Christensen, J. L., Wright, D. E., Wagers, A. J. & Weissman, I. L. Circulation and chemotaxis of fetal hematopoietic stem cells. PLoS Biol 2, E75 (2004). 10.1371/journal.pbio.0020075

63 Cain, T. L., Derecka, M. & McKinney-Freeman, S. The role of the haematopoietic stem cell niche in development and ageing. Nature Reviews Molecular Cell Biology (2024). 10.1038/s41580-024-00770-8

64 Mikkola, H. K. & Orkin, S. H. The journey of developing hematopoietic stem cells. Development 133, 3733–3744 (2006). 10.1242/dev.02568

65 Challen, G. A., Pietras, E. M., Wallscheid, N. C. & Signer, R. A. J. Simplified murine multipotent progenitor isolation scheme: Establishing a consensus approach for multipotent progenitor identification. Exp Hematol. 104, 55–63 (2021). 10.1016/j.exphem.2021.09.007

66 Akashi, K., Traver, D., Miyamoto, T. & Weissman, I. L. A clonogenic common myeloid progenitor that gives rise to all myeloid lineages. Nature 404, 193–197 (2000). 10.1038/35004599

67 Chen, M. Z. et al. A thiol probe for measuring unfolded protein load and proteostasis in cells. Nature Communications 8, 1–10 (2017). 10.1038/s41467-017-00203-5

68 Kim, W. et al. Systematic and quantitative assessment of the ubiquitin-modified proteome. Molecular Cell 44, 325–340 (2011). 10.1016/j.molcel.2011.08.025

69 Barton, B. M. et al. *IRE1α – XBP1 safeguards hematopoietic stem and progenitor cells by restricting pro-leukemogenic gene programs*. Vol. 26 (Springer US, 2025).

70 Lam, K. et al. The Proteostasis Network is a Therapeutic Target in Acute Myeloid Leukemia. Blood (2025). 10.1182/blood.2024026749

71 Jutzi, J. S. et al. CALR-mutated cells are vulnerable to combined inhibition of the proteasome and the endoplasmic reticulum stress response. Leukemia 37, 359–369 (2023). 10.1038/s41375-022-01781-0

72 Burgess, R. J., Zhao, Z., Nakada, D. & Morrison, S. J. Bmi1 suppresses protein synthesis and promotes proteostasis in hematopoietic stem cells. Genes Dev 36, 887–900 (2022). 10.1101/gad.349917.122

73 Li, Y. et al. Single-Cell Analysis of Neonatal HSC Ontogeny Reveals Gradual and Uncoordinated Transcriptional Reprogramming that Begins before Birth. Cell Stem Cell 27, 732–747.e737 (2020). 10.1016/j.stem.2020.08.001

74 Dikic, I. Proteasomal and Autophagic Degradation Systems. Annu Rev Biochem 86, 193–224 (2017). 10.1146/annurev-biochem-061516-044908

75 Gomez-Pastor, R., Burchfiel, E. T. & Thiele, D. J. Regulation of heat shock transcription factors and their roles in physiology and disease. Nature Reviews Molecular Cell Biology 19, 4–19 (2018). 10.1038/nrm.2017.73

76 Åkerfelt, M., Morimoto, R. I. & Sistonen, L. Heat shock factors: Integrators of cell stress, development and lifespan. Nature Reviews Molecular Cell Biology 11, 545–555 (2010). 10.1038/nrm2938

77 Kruta, M. et al. Hsf1 promotes hematopoietic stem cell fitness and proteostasis in response to ex vivo culture stress and aging. Cell Stem Cell 28, 1950–1965.e1956 (2021). 10.1016/j.stem.2021.07.009

78 Neef, D. W. et al. A direct regulatory interaction between chaperonin TRiC and stress-responsive transcription factor HSF1. Cell Reports 9, 955–966 (2014). 10.1016/j.celrep.2014.09.056

79 Shi, Y., Mosser, D. D. & Morimoto, R. I. Molecular chaperones as HSF1-specific transcriptional repressors. Genes and Development 12, 654–666 (1998). 10.1101/gad.12.5.654

80 Zou, J., Guo, Y., Guettouche, T., Smith, D. F. & Voellmy, R. Repression of heat shock transcription factor HSF1 activation by HSP90 (HSP90 complex) that forms a stress-sensitive complex with HSF1. Cell 94, 471–480 (1998). 10.1016/S0092-8674(00)81588-3

81 Anckar, J. & Sistonen, L. Regulation of HSF1 function in the heat stress response: Implications in aging and disease. Annual Review of Biochemistry 80, 1089–1115 (2011). 10.1146/annurev-biochem-060809-095203

82 Mendillo, M. L. et al. HSF1 drives a transcriptional program distinct from heat shock to support highly malignant human cancers. Cell 150, 549–562 (2012). 10.1016/j.cell.2012.06.031

83 Li, H. et al. The dynamics of hematopoiesis over the human lifespan. Nature Methods (2024). 10.1038/s41592-024-02495-0

84 Zhou, F. J. et al. Proteostasis Stress Drives Stem Cell Aging, Clonal Hematopoiesis and Leukemia. bioRxiv, 2025.2011.2027.690982 (2025). 10.1101/2025.11.27.690982

85 Hockemeyer, K. et al. The stress response regulator HSF1 modulates natural killer cell anti-tumour immunity. Nat Cell Biol 26, 1734–1744 (2024). 10.1038/s41556-024-01490-z

86 Yu, H. & Signer, R. Proteostasis Disruption in Inherited Bone Marrow Failure Syndromes. Blood (2025). 10.1182/blood.2024024956

87 Burwick, N., Coats, S. A., Nakamura, T. & Shimamura, A. Impaired ribosomal subunit association in Shwachman-Diamond syndrome. Blood 120, 5143–5152 (2012). 10.1182/blood-2012-04-420166

88 Sezgin, G. et al. Impaired growth, hematopoietic colony formation, and ribosome maturation in human cells depleted of Shwachman-Diamond syndrome protein SBDS. Pediatr Blood Cancer 60, 281–286 (2013). 10.1002/pbc.24300

89 Jaako, P. et al. eIF6 rebinding dynamically couples ribosome maturation and translation. Nature Communications 13, 1–11 (2022). 10.1038/s41467-022-29214-7

90 Moore Iv, J. B., Farrar, J. E., Arceci, R. J., Liu, J. M. & Ellis, S. R. Distinct ribosome maturation defects in yeast models of Diamond-Blackfan anemia and Shwachman-Diamond syndrome. Haematologica 95, 57–64 (2010). 10.3324/haematol.2009.012450

91 Garçon, L. et al. Ribosomal and hematopoietic defects in induced pluripotent stem cells derived from Diamond Blackfan anemia patients. Blood 122, 912–921 (2013). 10.1182/blood-2013-01-478321

92 Wong, C. C., Traynor, D., Basse, N., Kay, R. R. & Warren, A. J. Defective ribosome assembly in Shwachman-Diamond syndrome. Blood 118, 4305–4312 (2011). 10.1182/blood-2011-06-353938

93 Rujkijyanont, P., Adams, S. L., Beyene, J. & Dror, Y. Bone marrow cells from patients with Shwachman-Diamond syndrome abnormally express genes involved in ribosome biogenesis and RNA processing. British Journal of Haematology 145, 806–815 (2009). 10.1111/j.1365-2141.2009.07692.x

94 Ulirsch, J. C. et al. The Genetic Landscape of Diamond-Blackfan Anemia. American Journal of Human Genetics 103, 930–947 (2018). 10.1016/j.ajhg.2018.10.027

95 Draptchinskaia, N. et al. The gene encoding ribosomal protein S19 is mutated in Diamond-Blackfan anaemia. Nature Genetics 21, 169–175 (1999). 10.1038/5951

96 Finch, A. J. et al. Uncoupling of GTP hydrolysis from eIF6 release on the ribosome causes shwachman-diamond syndrome. Genes and Development 25, 917–929 (2011). 10.1101/gad.623011

97 Menne, T. F. et al. The Shwachman-Bodian-Diamond syndrome protein mediates translational activation of ribosomes in yeast. Nature Genetics 39, 486–495 (2007). 10.1038/ng1994

98 Calamita, P. et al. SBDS-Deficient Cells Have an Altered Homeostatic Equilibrium due to Translational Inefficiency Which Explains their Reduced Fitness and Provides a Logical Framework for Intervention. PLoS Genetics 13, 1–26 (2017). 10.1371/journal.pgen.1006552

99 Tan, S. et al. EFL1 mutations impair eIF6 release to cause Shwachman-Diamond syndrome. Blood 134, 277–290 (2019). 10.1182/blood.2018893404

100 In, K. et al. Shwachman-Bodian-Diamond syndrome (SBDS) protein deficiency impairs translation re-initiation from C/EBPα and C/EBPβ mRNAs. Nucleic Acids Research 44, 4134–4146 (2016). 10.1093/nar/gkw005

101 Bellanné-Chantelot, C. et al. Mutations in the SRP54 gene cause severe congenital neutropenia as well as Shwachman-Diamond – Like syndrome. Blood 132, 1318–1331 (2018). 10.1182/blood-2017-12-820308

102 Carapito, R. et al. Mutations in signal recognition particle SRP54 cause syndromic neutropenia with Shwachman-Diamond-like features. Journal of Clinical Investigation 127, 4090–4103 (2017).

103 Nanua, S. et al. Activation of the unfolded protein response is associated with impaired granulopoiesis in transgenic mice expressing mutant Elane. Blood 117, 3539–3547 (2011). 10.1182/blood-2010-10-311704

104 Grenda, D. S. et al. Mutations of the ELA2 gene found in patients with severe congenital neutropenia induce the unfolded protein response and cellular apoptosis. Blood 110, 4179–4187 (2007). 10.1182/blood-2006-11-057299

105 Köllner, I. et al. Mutations in neutrophil elastase causing congenital neutropenia lead to cytoplasmic protein accumulation and induction of the unfolded protein response. Blood 108, 493–500 (2006). 10.1182/blood-2005-11-4689

106 Massullo, P. et al. Aberrant subcellular targeting of the G185R neutrophil elastase mutant associated with severe congenital neutropenia induces premature apoptosis of differentiating promyelocytes. Blood 105, 3397–3404 (2005). 10.1182/blood-2004-07-2618

107 Wiesmeier, M., Gautam, S., Kirschnek, S. & Häcker, G. Characterisation of neutropenia-associated neutrophil elastase mutations in a murine differentiation model in vitro and in vivo. PLoS ONE 11, 1–20 (2016). 10.1371/journal.pone.0168055

108 Nustede, R. et al. ELANE mutant-specific activation of different UPR pathways in congenital neutropenia. British Journal of Haematology 172, 219–227 (2016). 10.1111/bjh.13823

109 Doll, L., Welte, K., Skokowa, J. & Bajoghli, B. A JAGN1-associated severe congenital neutropenia zebrafish model revealed an altered G-CSFR signaling and UPR activation. Blood Advances (2024).

110 Van Nieuwenhove, E. et al. Defective Sec61α1 underlies a novel cause of autosomal dominant severe congenital neutropenia. Journal of Allergy and Clinical Immunology 146, 1180–1193 (2020). 10.1016/j.jaci.2020.03.034

111 Schürch, C. et al. SRP54 mutations induce congenital neutropenia via dominant-negative effects on XBP1 splicing. Blood 137, 1340–1352 (2021). 10.1182/blood.2020008115

112 Robertson, N. et al. A disease-linked lncRNA mutation in RNase MRP inhibits ribosome synthesis. Nature Communications 13 (2022). 10.1038/s41467-022-28295-8

113 Zhang, F. et al. Human SAMD9 is a poxvirus-activatable anticodon nuclease inhibiting codon-specific protein synthesis. Science Advances 9 (2023). 10.1126/sciadv.adh8502

114 Thomas, M. E. et al. Pediatric MDS and bone marrow failure-associated germline mutations in SAMD9 and SAMD9L impair multiple pathways in primary hematopoietic cells. Leukemia 35, 3232–3244 (2021). 10.1038/s41375-021-01212-6

115 Cai, X. et al. Runx1 Deficiency Decreases Ribosome Biogenesis and Confers Stress Resistance to Hematopoietic Stem and Progenitor Cells. Cell Stem Cell 17, 165–177 (2015). 10.1016/j.stem.2015.06.002

116 Williams, S. A., Longerich, S., Sung, P., Vaziri, C. & Kupfer, G. M. The E3 ubiquitin ligase RAD18 regulates ubiquitylation and chromatin loading of FANCD2 and FANCI. Blood 117, 5078–5087 (2011). 10.1182/blood-2010-10-311761

117 Parmar, K. et al. Hematopoietic stem cell defects in mice with deficiency of Fancd2 or Usp1. Stem Cells 28, 1186–1195 (2010). 10.1002/stem.437

118 Gueiderikh, A. et al. Fanconi anemia A protein participates in nucleolar homeostasis maintenance and ribosome biogenesis. Science Advances 7, 1–12 (2021). 10.1126/sciadv.abb5414

119 Oda, T., Hayano, T., Miyaso, H., Takahashi, N. & Yamashita, T. Hsp90 regulates the Fanconi anemia DNA damage response pathway. Blood 109, 5016–5026 (2007). 10.1182/blood-2006-08-038638

120 Karras, G. I. et al. HSP90 Shapes the Consequences of Human Genetic Variation. Cell 168, 856–866.e812 (2017). 10.1016/j.cell.2017.01.023

121 Sondalle, S. B., Longerich, S., Ogawa, L. M., Sung, P. & Baserga, S. J. Fanconi anemia protein FANCI functions in ribosome biogenesis. Proceedings of the National Academy of Sciences of the United States of America 116, 2561–2570 (2019). 10.1073/pnas.1811557116

122 Li, L., Wang, Z. V., Hill, J. A. & Lin, F. New autophagy reporter mice reveal dynamics of proximal tubular autophagy. Journal of the American Society of Nephrology 25, 305–315 (2014). 10.1681/ASN.2013040374

123 Georgiades, P. et al. vavCre transgenic mice: A tool for mutagenesis in hematopoietic and endothelial lineages. Genesis (United States) 34, 251–256 (2002). 10.1002/gene.10161

124 Le Masson, F. et al. Identification of heat shock factor 1 molecular and cellular targets during embryonic and adult female meiosis. Mol Cell Biol 31, 3410–3423 (2011). 10.1128/MCB.05237-11

125 Oguro, H., Ding, L. & Morrison, Sean J. SLAM Family Markers Resolve Functionally Distinct Subpopulations of Hematopoietic Stem Cells and Multipotent Progenitors. Cell Stem Cell 13, 102–116 (2013). 10.1016/j.stem.2013.05.014

126 Pietras, Eric M. et al. Functionally Distinct Subsets of Lineage-Biased Multipotent Progenitors Control Blood Production in Normal and Regenerative Conditions. Cell Stem Cell 17, 246 (2015). 10.1016/j.stem.2015.07.003

127 Hardy, R. R., Carmack, C. E., Shinton, S. A., Kemp, J. D. & Hayakawa, K. Resolution and characterization of pro-B and pre-pro-B cell stages in normal mouse bone marrow. Journal of Experimental Medicine 173, 1213–1225 (1991). 10.1084/jem.173.5.1213

128 Picelli, S. et al. Full-length RNA-seq from single cells using Smart-seq2. Nature Protocols 9, 171–181 (2014). 10.1038/nprot.2014.006

129 Dobin, A. et al. STAR: ultrafast universal RNA-seq aligner. Bioinformatics 29, 15–21 (2013). 10.1093/bioinformatics/bts635

130 Schmieder, R. & Edwards, R. Quality control and preprocessing of metagenomic datasets. Bioinformatics 27, 863–864 (2011). 10.1093/bioinformatics/btr026

131 Li, H. et al. The Sequence Alignment/Map format and SAMtools. Bioinformatics 25, 2078–2079 (2009). 10.1093/bioinformatics/btp352

132 Anders, S., Pyl, P. T. & Huber, W. HTSeq—a Python framework to work with high-throughput sequencing data. Bioinformatics 31, 166–169 (2015). 10.1093/bioinformatics/btu638

133 Love, M. I., Huber, W. & Anders, S. Moderated estimation of fold change and dispersion for RNA-seq data with DESeq2. Genome Biology 15, 550 (2014). 10.1186/s13059-014-0550-8

134 Subramanian, A. et al. Gene set enrichment analysis: A knowledge-based approach for interpreting genome-wide expression profiles. Proceedings of the National Academy of Sciences 102, 15545–15550 (2005). 10.1073/pnas.0506580102

135 Liberzon, A. et al. Molecular signatures database (MSigDB) 3.0. Bioinformatics 27, 1739–1740 (2011). 10.1093/bioinformatics/btr260

136 Liberzon, A. et al. The Molecular Signatures Database (MSigDB) hallmark gene set collection. Cell Systems 1(**6**), 417–425 (2015). 10.1016/j.cels.2015.12.004

137 Ashburner, M. et al. Gene Ontology: tool for the unification of biology. Nature Genetics 25, 25–29 (2000). 10.1038/75556

138 Hänzelmann, S., Castelo, R. & Guinney, J. GSVA: gene set variation analysis for microarray and RNA-seq data. BMC Bioinformatics 14, 7 (2013). 10.1186/1471-2105-14-7

139 Fang, Z., Liu, X. & Peltz, G. GSEApy: a comprehensive package for performing gene set enrichment analysis in Python. Bioinformatics 39 (2023). 10.1093/bioinformatics/btac757

