## Supplemental Material for "Proteostasis Remodeling Across Development Defines Fetal, Neonatal, and Adult Hematopoietic Stem Cell States"

### Supplementary Figure S1

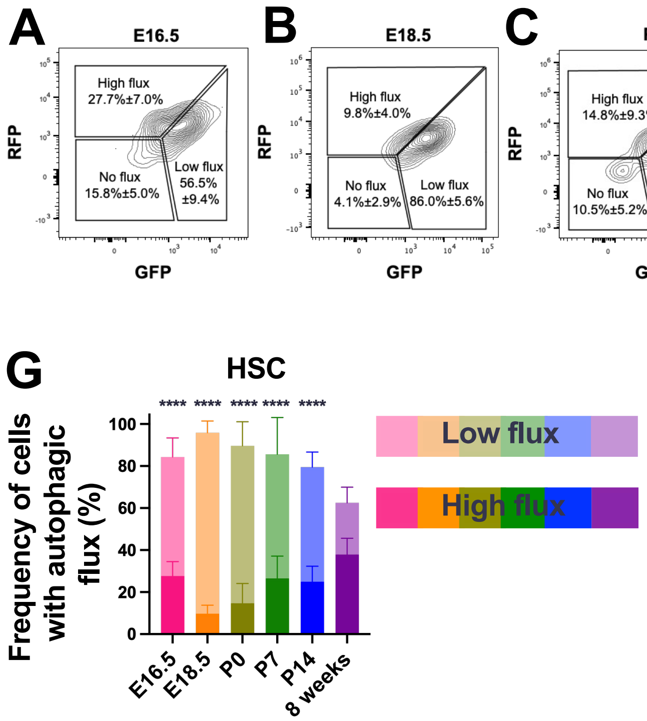

**Supplementary Figure S1**

(A-F) Representative flow cytometry plots and frequency of HSC populations with high, low, and no autophagic flux based on RFP and GFP expression in CAG-RFP-EGFP-LC3 (A) E16.5, (B) E18.5, (C) P0, (D) P7, (E) P14, and (F) 8-week-old mice.

(G) Frequency of HSCs with autophagic flux as measured by the combined population of $RFP^{+}EGFP^{-}$ (high autophagic flux) and $RFP^{+}EGFP^{+}$ (low autophagic flux) cells at E16.5, E18.5, P0, P7, and P14 (n = 8-21 mice/age in 2-5 experiments).

Data represent mean ± standard deviation. Statistical significance was assessed using a one-way ANOVA followed by Dunnett's multiple comparisons test relative to the 8-week-old timepoint. ****P<0.0001.

### Supplementary Figure S2

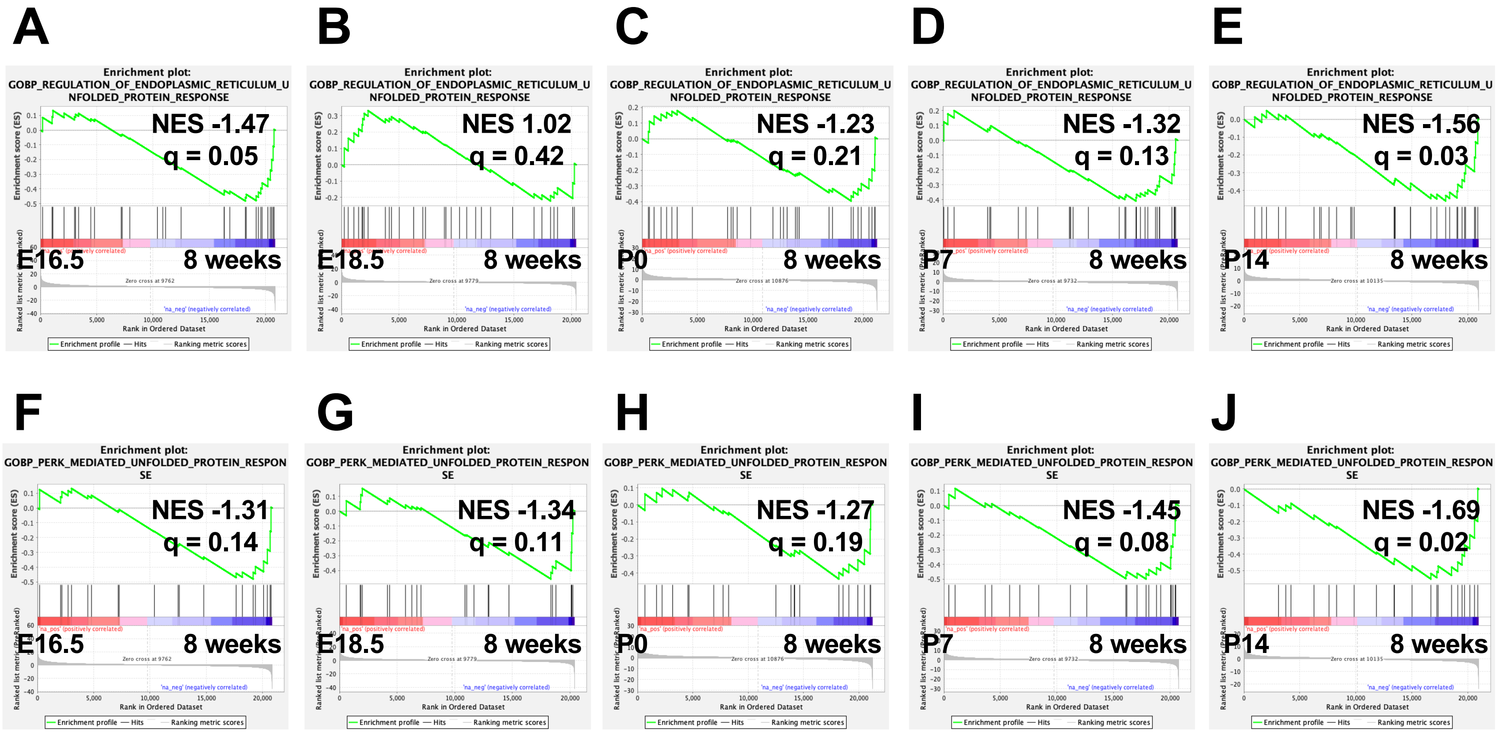

**Supplementary Figure S2**

(A-E) Enrichment plots for the “Regulation of endoplasmic reticulum unfolded protein response” gene set based on RNA-sequencing data in HSCs (GSE128759) from E16.5 (A), E18.5 (B), P0 (C), P7 (D), and P14 (E) mice compared to 8-week-old mice.

(F-J) Enrichment plots for the “Regulation of PERK mediated unfolded protein response” gene set based on RNA-sequencing data in HSCs (GSE128759) from E16.5 (F), E18.5 (G), P0 (H), P7 (I), and P14 (J) mice compared to 8-week-old mice.

Normalized enrichment scores (NES) and false discovery rates (q) are shown.

### Supplementary Figure S3

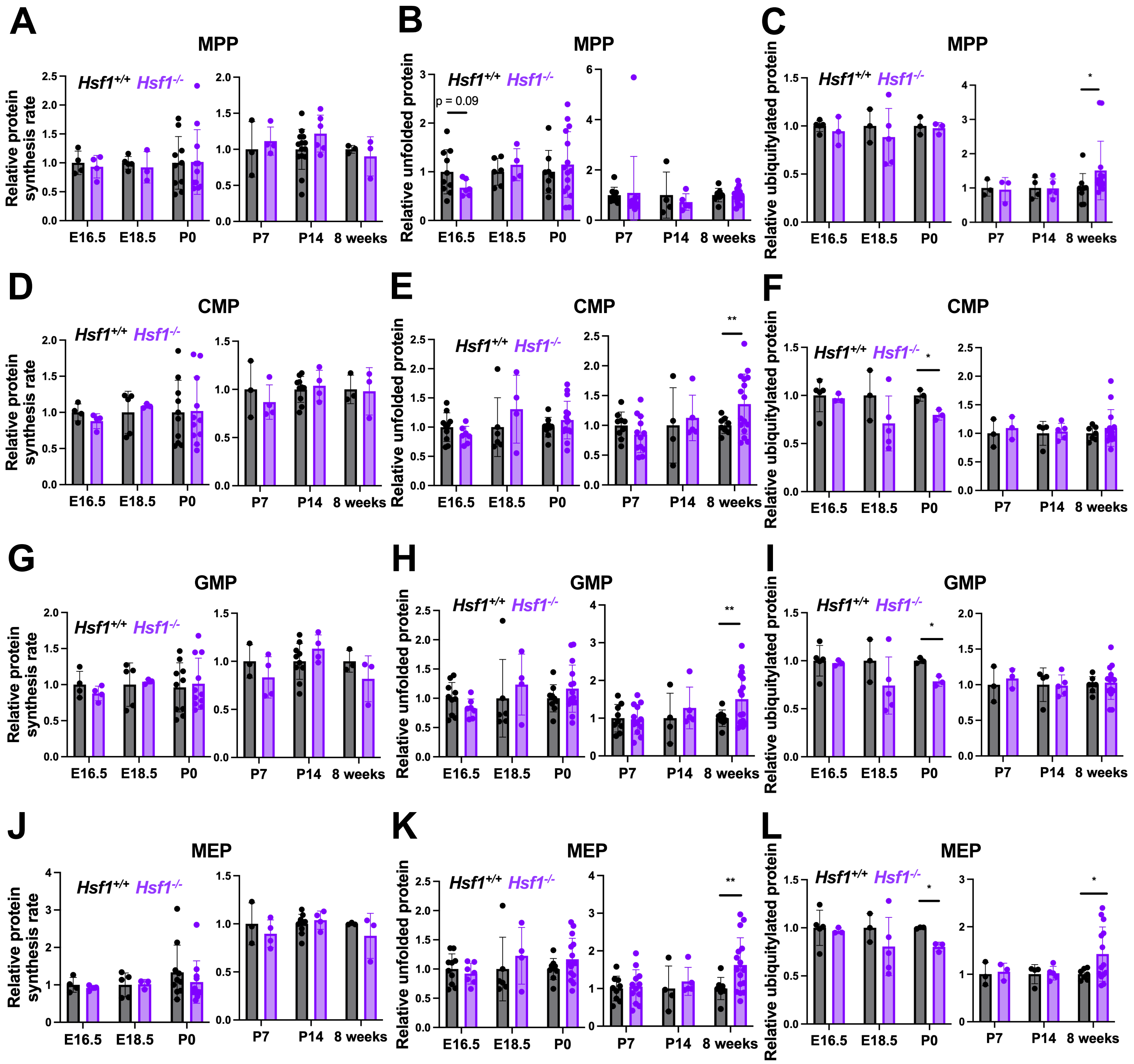

**Supplementary Figure S3**

(A, D, G, J) Relative protein synthesis rates in liver (left) and bone marrow (right) MPPs (A), CMPs (D), GMPs (G), and MEPs (J) from *Hsf1*$^{+/+}$ and *Hsf1*$^{-/-}$ mice at E16.5, E18.5, P0, P7, P14, and 8 weeks. OP-Puro MFI is normalized to *Hsf1*$^{+/+}$ HSCs at each respective age (n = 3-14 mice/genotype/age).

(B, E, H, K) Quantity of unfolded protein in liver (left) and bone marrow (right) MPPs (B), CMPs (E), GMPs (H), and MEPs (K) from *Hsf1*$^{+/+}$ and *Hsf1*$^{-/-}$ mice at E16.5, E18.5, P0, P7, P14, and 8 weeks. TMI MFI is normalized to *Hsf1*$^{+/+}$ HSCs at each respective age (n = 4-16 mice/genotype/age/age).

(C, F, I, L) Relative quantity of ubiquitylated protein in liver (left) and bone marrow (right) MPPs (C), CMPs (F), GMPs (I), and MEPs (L) from *Hsf1*$^{+/+}$ and *Hsf1*$^{-/-}$ mice at E16.5, E18.5, P0, P7, P14, and 8 weeks. MFI is normalized to *Hsf1*$^{+/+}$ HSCs at each respective age (n = 3-14 mice/genotype/age).

Data represent mean ± standard deviation. Differences between groups were assessed by Welch's t-test. *P<0.05; **P<0.01.

### Supplementary Figure S4

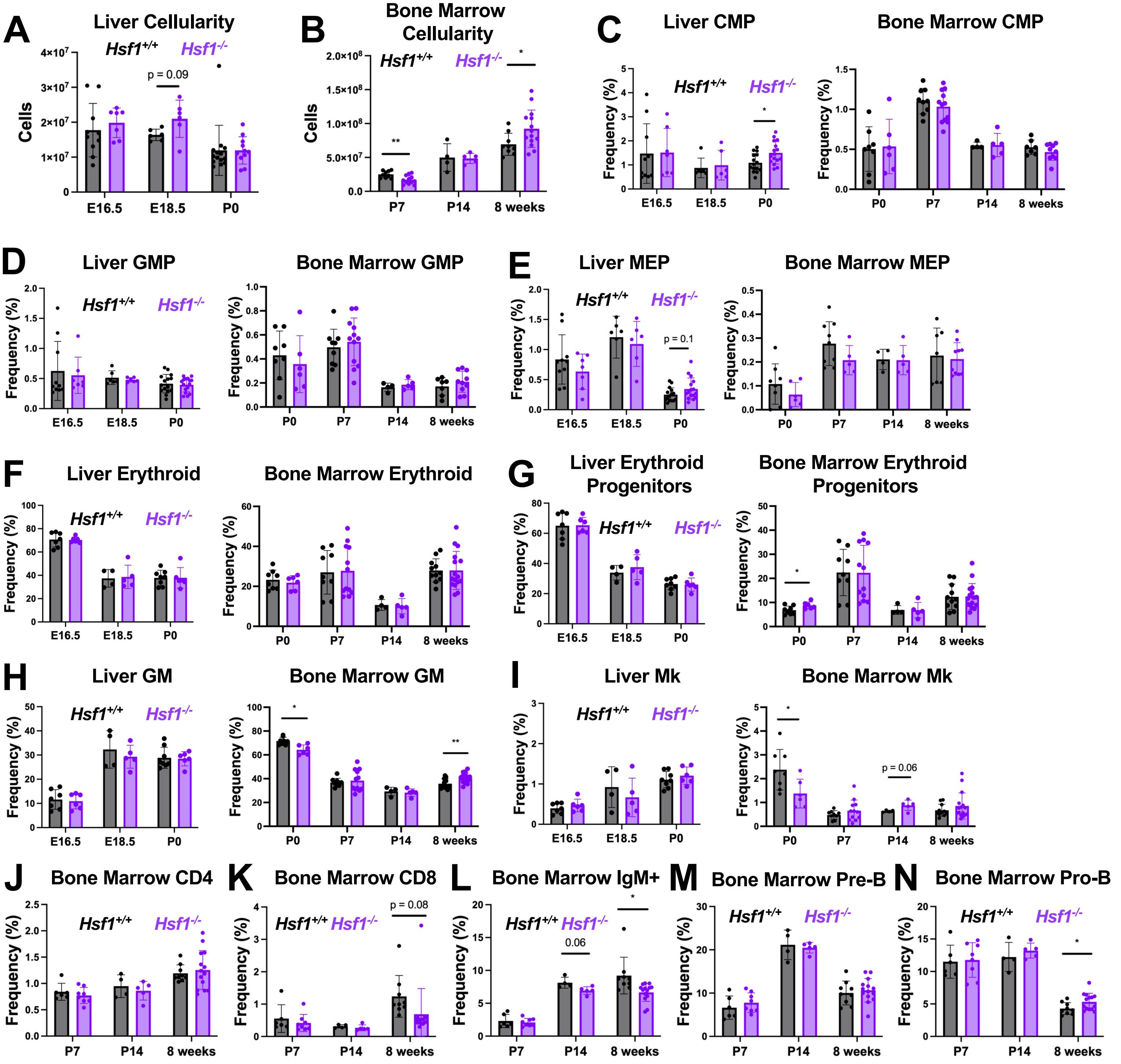

**Supplementary Figure S4**

(A-B) Liver (A) and bone marrow (B) cellularity in *Hsf1*$^{+/+}$ and *Hsf1*$^{-/-}$ mice at E16.5, E18.5, P0, P7, P14, and 8 weeks (n = 4-14 mice/genotype/age).

(C-I) Frequency of CMP (C), GMP (D), MEP (E), Ter119$^{+}$ erythroid (F), CD71$^{+}$Ter119$^{+}$ erythroid progenitor (G), Gr1$^{+}$CD11b$^{+}$ GM (H), and CD41$^{+}$ megakaryocyte (I) cell populations in *Hsf1*$^{+/+}$ and *Hsf1*$^{-/-}$ mice at E16.5, E18.5, P0, P7, P14, and 8 weeks (n = 4-16 mice/genotype/age).

(J-N) Frequency of bone marrow CD4$^{+}$ (J), CD8$^{+}$ (K), B220$^{+}$IgM$^{+}$ B (L), B220$^{+}$IgM$^{-}$CD43$^{-}$ Pre-B (M), and B220$^{+}$IgM$^{-}$CD43$^{+}$ Pro-B (N) cell populations at P7, P14, and 8 weeks in *Hsf1*$^{+/+}$ and *Hsf1*$^{-/-}$ mice (n = 4-14 mice/genotype/age).

Data represent mean ± standard deviation. Differences between groups were assessed by Welch's t-test. *P<0.05; **P<0.01.
